# A fast and versatile method for simultaneous HCR, immunohistochemistry and EdU labeling (SHInE)

**DOI:** 10.1101/2022.10.13.512152

**Authors:** Aida Ćorić, Alexander W. Stockinger, Petra Schaffer, Dunja Rokvić, Kristin Tessmar-Raible, Florian Raible

**Author notes:** equal contribution (Aida Ćorić, Alexander W. Stockinger).

## Abstract

Access to newer, fast and cheap sequencing techniques, particularly on the single-cell level, have made transcriptomic data of tissues or single cells accessible to many researchers. As a consequence, there is increased need for *in situ* visualization of gene expression or encoded proteins to validate, localize or help interpret such sequencing data, as well as put them in context with cellular proliferation. A particular challenge for labeling and imaging transcripts are complex tissues that are often opaque and/or pigmented, preventing easy visual inspection. Here we introduce a versatile protocol that combines *in situ* hybridization chain reaction (HCR), immunohistochemistry (IHC) and proliferative cell labeling using 5-ethynyl-2’-deoxyuridine (EdU), and demonstrate its compatibility with tissue clearing. As a proof-of-concept, we show that our protocol allows for the parallel analysis of cell proliferation, gene expression and protein localization in bristleworm heads and trunks.

## Introduction

Advancements in sequencing techniques have significantly increased the number of organisms for which molecular research can be conducted in. Moreover, single-cell RNA sequencing allows such investigations on the level of individual cell types (Fonseca et al., 2016; Stark et al., 2019). In turn, these high throughput methods have generated a need for validating digital expression data in the corresponding organism, specifically for the visualization of gene-expression on the three-dimensional level.

In recent years, strategies complementary to traditional enzyme-coupled *in situ* hybridization (ISH) have been developed. One of them is *in situ* hybridization chain reaction (HCR). *In situ* HCR is faster to perform than traditional ISH, works at lower temperatures which aids maintaining tissue integrity, and can be multiplexed by using different fluorophores compatible with fluorescent imaging (Choi et al., 2016), making *in situ* HCR an attractive companion technique to single cell sequencing.

For *in situ* HCR, short DNA probes are synthesized that are complementary to the target transcript. These probes can be designed using web-tools and ordered on bulk scale, similar to primers, which makes production fast and affordable. Amplifier oligonucleotides carrying fluorophores are then annealed to an overhang region on the probes, triggering the release of a hairpin structure, and thereby revealing the binding sequence for another amplifier molecule. This leads to a chain reaction of linear signal amplification. The latest version of the method, termed HCR 3.0 (Choi et al., 2018), employs split initiator sequences: Two DNA probes form a probe pair and have to successfully bind next to each other to form a shared initiator sequence. Off-target binding or trapping of individual probes therefore should be less likely to cause signal, improving sensitivity of the method.

Whereas high sensitivity is an important feature for the use of *in situ* HCR to validate scRNAseq data, there are also challenges and open questions to the use of this technique: First, the acquisition of images from *in-situ*-HCR-stained samples by confocal microscopy bears potential problems in larger specimens, as opaqueness of tissue or pigmentation are known to prevent deep imaging (Tainaka et al., 2015). While various tissue clearing methods have been developed to overcome imaging problems in such specimens (Vieites-Prado and Renier, 2021), their compatibility with RNA detection has not been systematically assessed.

Secondly, while visualizing RNA is a relevant experimental aim, there are contexts where co-visualisation of proteins by immunohistochemistry (IHC) would yield additional insights. For example, such co-detection could help to benchmark gene expression in cells or tissues for which protein markers are already established, or to directly assess ratios of RNA and corresponding protein (Albayrak et al., 2016; Darmanis et al., 2015). Combining the visualization of protein and RNA can therefore be used to study developmental processes or gene regulation in ways each technology on its own would not allow.

Few protocols have been published that combine the convenience, flexibility and sensitivity of *in situ* HCR with immunostainings. Those describing a combined approach either rely on the availability of commercial antibodies and proprietary technology (Schwarzkopf et al., 2021) or follow a longer protocol that performs immunostaining after finishing *in situ* HCR (Elagoz et al., 2022; Ibarra-García-Padilla et al., 2021), which delays workflows and potentially lowers the achieved signal.

Here we present a protocol (termed Simultaneous HCR, Immunohistochemistry and EdU labeling / SHInE; Figure 1) that addresses both of these challenges: SHInE combines *in situ* HCR with IHC, allowing co-detection of RNA and protein. By introducing the primary antibody incubation of the IHC simultaneously to the amplification step of *in situ* HCR, SHInE saves experimental time and keeps the number of washing steps following amplification to a minimum. Moreover, SHInE is compatible with tissue clearing using the recently developed DEEP-Clear method (Pende et al., 2020), extending the portfolio of labeling techniques for this method.

**Figure 1.**
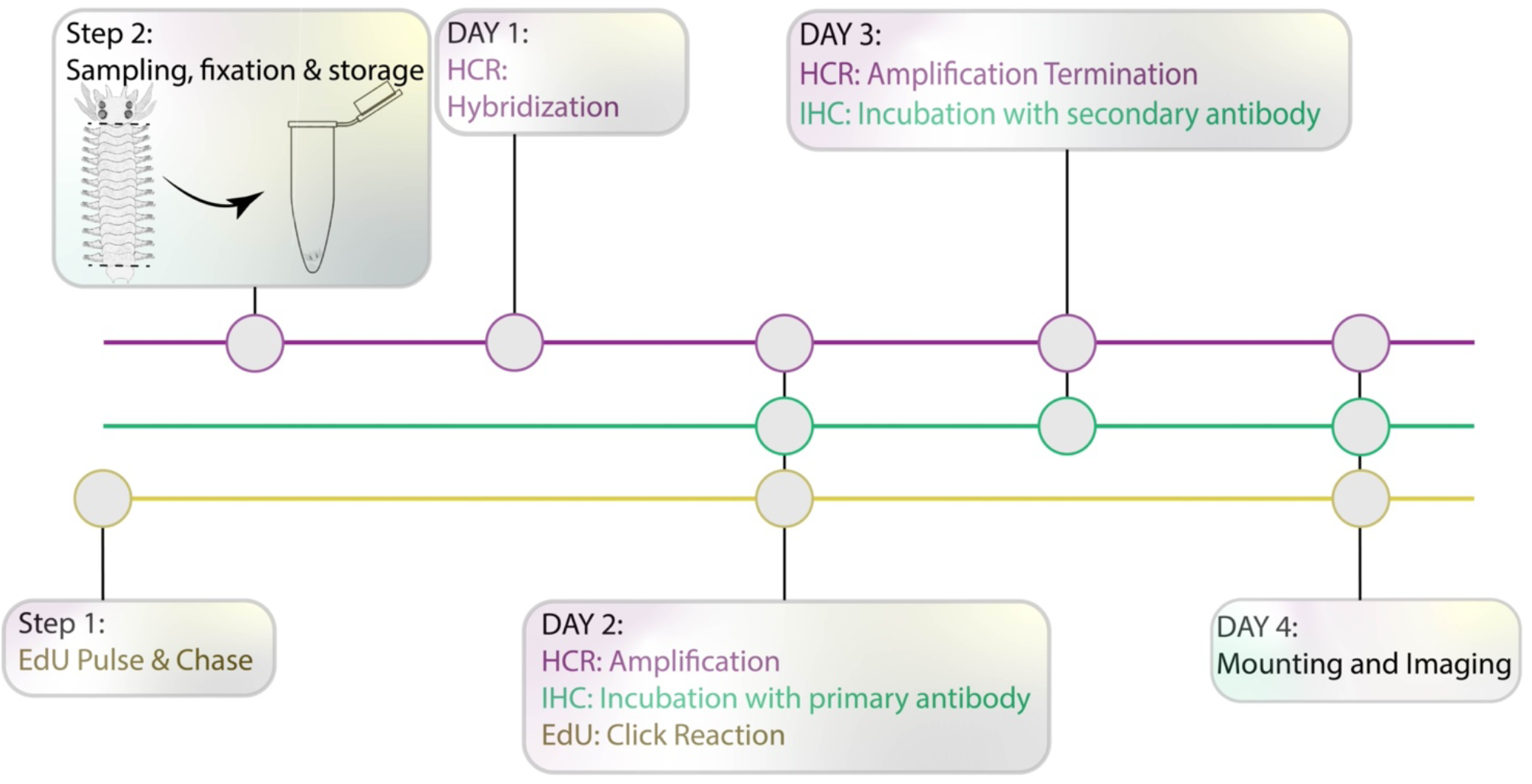
Experimental setup for simultaneous in situ HCR, Immunohistochemistry and EdU labeling. Following the initial treatment with EdU, animals are sampled and fixed for in situ hybridization chain reaction (HCR). Our protocol allows for the implementation of HCR amplification, EdU detection (click reaction) and incubation with primary antibody for IHC on Day 2, resulting in a total of 4 days from sampling to imaging.

Our protocol also offers the flexibility of using non-commercial, self-designed HCR probes along with custom antibodies and home-made buffers. In addition, SHInE has been adapted to include additional molecular assays, such as the labeling of proliferating cells with 5-ethynyl-2’-deoxyuridine (EdU) (Salic and Mitchison, 2008; Zattara and Özpolat, 2020) further enabling the study of complex biological processes.

To demonstrate the functionality of our protocol, we provide results for different sets of HCR probes, different antibodies, and in different tissues of the marine bristle worm *Platynereis dumerilii*. This invertebrate has gained importance as a model organism in various biological fields, including evolution, development, regeneration, neurobiology, reproduction, chronobiology and ecology (reviewed in (Özpolat et al., 2021a)). As in other model systems, conventional, riboprobe-based *in situ* hybridization has been successfully established for gene expression studies in this species (Tessmar-Raible et al., 2005), SHInE provides a new and flexible tool for studying a broad range of biological processes in *Platynereis dumerilii*, and likely can be adapted to other species.

## Materials and Methods

A schematized workflow of the SHInE protocol is presented in Figure 1. A detailed version of the protocol, including all reagents and buffers, has been deposited on protocols.io (DOI: dx.doi.org/10.17504/protocols.io.5qpvobnyzl4o/v1). This version contains detailed steps with estimated time requirements, pause points, and steps that can be adjusted according to specific experimental needs (Supplementary File 1).

### Animal husbandry

*Platynereis dumerilii* worms were kept in continuous culture at the Max Perutz Labs Vienna Marine Facility under a 16:8 light:dark regime. For details on animal culture, see (Hauenschild and Fischer, 1969; Kuehn et al., 2019).

### Posterior amputations

Sibling worms aged 3-5 months and of a size between 40 and 50 segments were sampled. Trunk pieces were surgically amputated by anesthetizing animals in 7,5% MgCl2 mixed 1:1 with artificial sea water, then removing all segments posterior to segment nr. 30 with a clean scalpel cut perpendicular to the animals’ body axis, similar to previous reports (Planques et al., 2018)

### Sampling of worm heads for *in situ* HCR

Wild-type worms were sampled at zeitgeber time (ZT) 22.

### EdU pulse labeling & click reaction

Animals were transferred into glass beakers and incubated with 10 μM EdU in NSW/ASW (mix 1:1) at the onset of darkness (ZT16) for 6 hours. Worm heads were subsequently sampled at ZT22.

For EdU labeling of worm blastemas, worms were put into a glass beaker and incubated with 10 μM EdU in ASW for 1h. No chase was performed for these experiments.

The click reaction was performed according to the manufacturer’s protocol using the Click-iT™ EdU Cell Proliferation Kit for Imaging, Alexa Fluor™ 488 dye.

### Microscopy parameters & Image processing

Images were taken on a Zeiss LSM 700 inverse confocal microscope using Plan-Apochromat 20x/0.8, WD 0.55 mm and LD LCI Plan-Apochromat 25x/0.8 mm (oil immersion) lenses. Contrast was adjusted in FIJI (Schindelin et al., 2012) and panels were arranged in Adobe Illustrator.

### HCR Probes

HCR probes were designed against the listed target genes (**table 1**) using a probe maker tool developed by Ryan Null (Kuehn et al., 2022) that was installed using Python v3.9.2 and jupyterlab 3.0.10.

**Table 1.** Target genes, amplifiers and used fluorophores for the HCR probes reported in this study. Respective probe sequences can be found in **Supplementary File 2**.

| **Gene** | **Reference / Source** | **Amplifier** | **Fluorophore** |
| --- | --- | --- | --- |
| *Pladu_hox3* | GenBank JQ424894.1 | B2 | Alexa 647 |
| *Pladu_pdp1* | this study (GenBank ID pending) | B1 | Alexa 546 |
| *Pladu_per* | this study (GenBank ID pending) | B2 | Alexa 647 |
| *Subdo_myc* | this study (GenBank ID pending) | B2 | Alexa 647 |

The number of probe pairs depended on the length of each target sequence. As detailed in **Supplementary File 2**, we used 30 probe pairs for *Platynereis hox3*, 49 probe pairs for *Platynereis pdp1*, and 27 probe pairs for *Platynereis per*.

### Antibodies

**Table 2.**
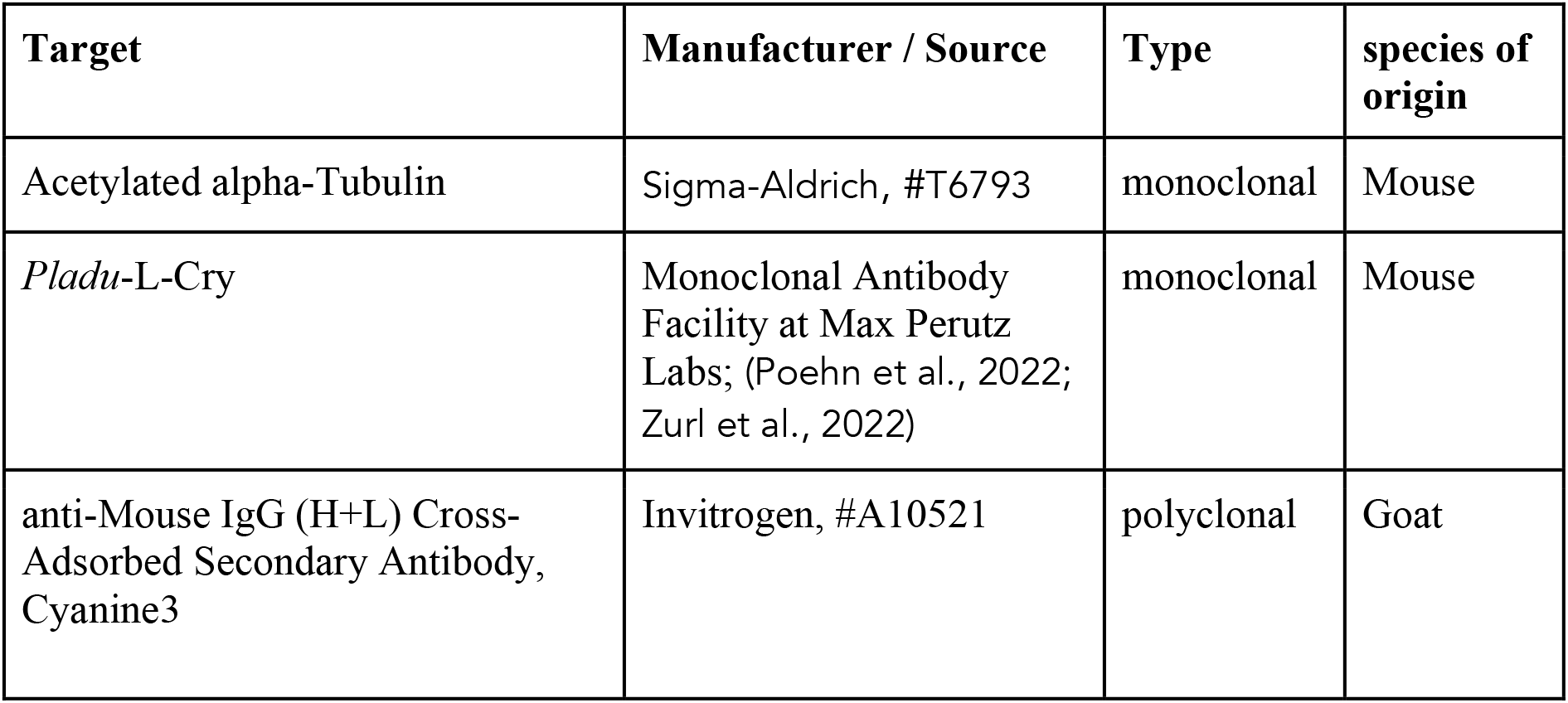
Target proteins and specifications of the antibodies used in this study.

| **Target** | **Manufacturer / Source** | **Type** | **species of origin** |
| --- | --- | --- | --- |
| Acetylated alpha-Tubulin | Sigma-Aldrich, #T6793 | monoclonal | Mouse |
| *Pladu*-L-Cry | Monoclonal Antibody Facility at Max Perutz Labs; (Poehn et al., 2022; Zurl et al., 2022) | monoclonal | Mouse |
| anti-Mouse IgG (H+L) Cross-Adsorbed Secondary Antibody, Cyanine3 | Invitrogen, #A10521 | polyclonal | Goat |

## Results

### SHInE allows co-detection of HCR probes, protein and nuclear labels in whole-mount samples

To address the ability of SHInE to co-visualize RNA, protein and S-phase-labeled nuclei, we produced and imaged two sets of samples. On the one hand, we used *Platynereis* heads to correlate the expression of the circadian clock gene *period* with the photoreceptor protein L-Cryptochrome (L-Cry) and the proliferation marker EdU (Figure 2). On the other hand, we assessed expression of the posterior stem cell marker *hox3* with stabilized microtubules and EdU in regenerating tail samples (Figure 3). In both cases, background staining in the respective channels was controlled by the use of HCR probes directed against a gene of the sponge *Suberites domuncula*. Moreover, obtained patterns were compared against the result of published expression studies to validate the accuracy of our method.

**Figure 2.**
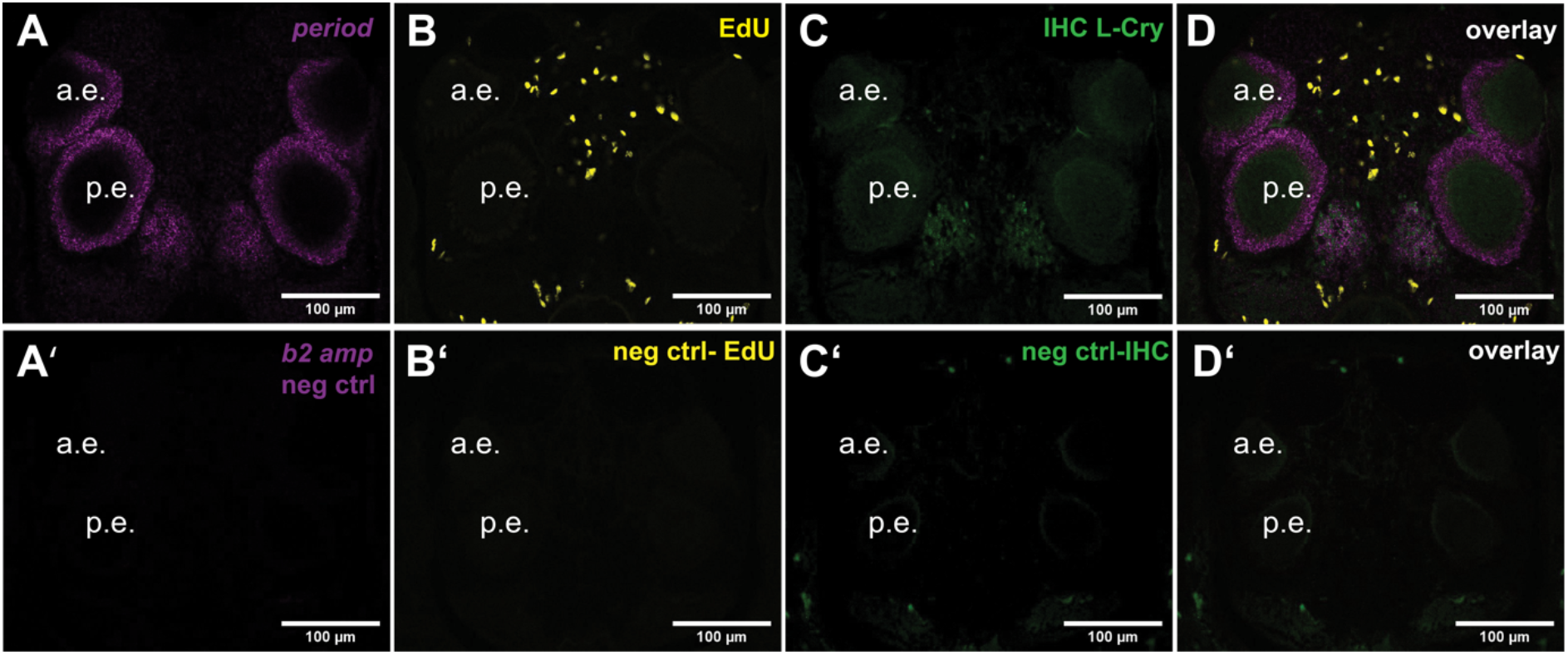
Co-detection of HCR probes, immunolabels and EdU in Platynereis heads. Tissues were processed according to the SHInE protocol and imaged on a confocal microscope. Platynereis heads show staining for period in both pairs of eyes and in an oval-shaped domain of the posterior brain (A). Sparse proliferation of cells can be observed throughout the worm head, but not in the eyes or the oval-shaped posterior domain (B). L-Cry, a photoreceptor protein, is found in the eyes and the oval-shaped posterior domain (C). The overlay (D) shows strong co-localization of L-Cry and period, while the respective negative controls (A’-D’) exhibit very low background signal. a. e.: anterior eye; p.e.: posterior eye.

**Figure 3.**
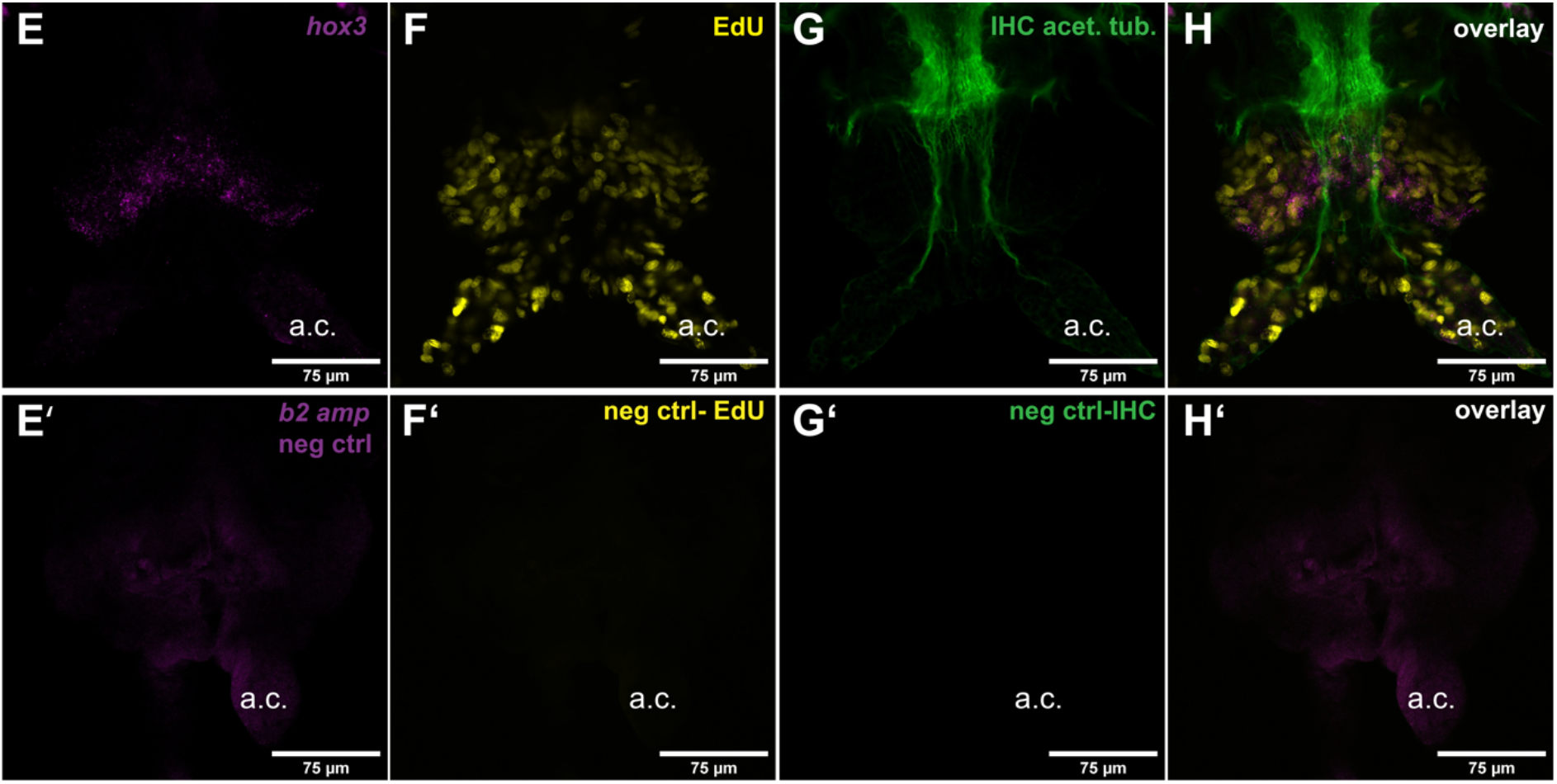
Co-detection of HCR probes, immunolabels and EdU in posterior regenerates. Posterior regenerating tissue at three days post caudal amputation contains a well-defined area of hox3 expression, referred to as segment addition zone (E). Proliferating cells can be observed throughout the regenerating tissue, especially in the anal cirri and the area surrounding the segment addition zone (F). The acetylated tubulin staining helps orientation within the tissue and shows the ventral nerve cord and its projections into the anal cirri (G). As in Figure 2, the respective negative controls (E’-H’) exhibit little to no background signal; some autofluorescence can be observed in the HCR negative control (E’), but lacks the granularity and intensity of HCR signal and can therefore be easily distinguished; a. c.: anal cirrus.

### Co-visualisation of L-Cry, *period* mRNA and EdU in heads

One of the interesting aspects about *Platynereis dumerilii* is its use as a functional molecular model system for chronobiological analyses (Özpolat et al., 2021b; Raible and Tessmar-Raible, 2014). At least two endogenous timing systems – a ~24hr (plastic circadian-circalunidian) and a monthly (circalunar) one – co-exist in *Platynereis* and are influenced by ambient light conditions (Zantke et al., 2013) (Poehn et al., 2022; Zurl et al., 2022). *Period* is a key gene of the circadian clock (Glossop and Hardin, 2002). The *Platynereis period* ortholog has previously been shown to be expressed in the head using riboprobe-based whole-mount *in situ* hybridization (Zantke et al., 2013). In accordance with the published pattern, HCR probes designed against *Platynereis period* demarcate cells in the posterior medial brain as well as in the eyes (Figure 2A,D). In addition to this established pattern, we noted weaker, more ubiquitous *period* staining, which cannot be seen in the negative control (cf. arrowheads in Figure 4B/B’). Whereas this expression is not seen in the conventional *in situ* hybridization for *period*, it might be attributed to the higher sensitivity of HCR and its ability to detect single mRNA molecules (Shah et al., 2016).

**Figure 4.**
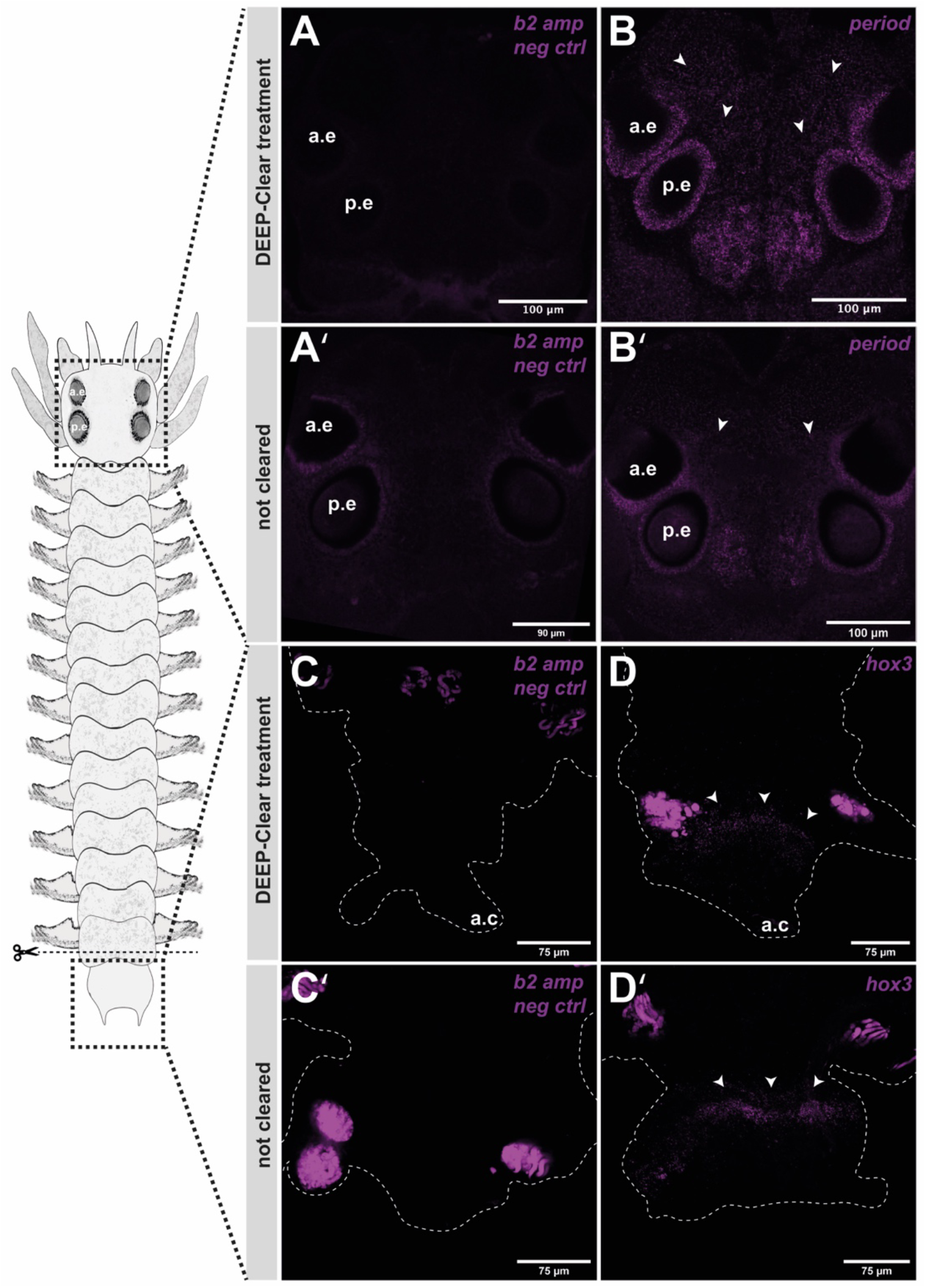
Compatibility of HCR detection with tissue clearing Platynereis heads show differences in background signal in cleared vs. not-cleared individuals. Autofluorescence can be seen around and in the eyes in the negative control for untreated samples, while there is no such signal in the negative control of cleared heads (A-A’). Pladu-period signal is strong around the eyes and in the nuclei of the posterior medial brain in P.dumerilii. Stronger HCR intensity is observed following tissue clearing, which highlights additional, less centralized signal of Pladu-period (B-B’). HCR signal intensity is less affected by clearing in Platynereis blastema. Autofluorescence is minimal in the negative control in untreated, as well as treated samples (C-C’). Pladu-hox3 signal is clearly defined and restricted to the segment addition zone (D-D’). Enhancement of signal intensity is not observed following tissue clearing.

In order to co-assess cell proliferation in the head during the night, we incubated worms in EdU solution from darkness onset to 2 hours before the lights turned (i.e., for a total of 6 hours). Visualization of EdU using click chemistry revealed positive cells in the anterior part of the head, in a dispersed, salt-and-pepper pattern with few cells also localizing to the posterior of the head (Figure 2B,D). At the investigated stage, we did not observe many proliferating cells in the eyes, which exhibit continuous growth and noticeably increase in size before sexual maturation (Fischer et al., 2010; Özpolat et al., 2021a; Pende et al., 2020).

To test the compatibility of SHInE with IHC, immunostainings were performed on the same samples using antibodies raised against *Platynereis* L-Cryptochrome (L-Cry). L-Cry is a photoreceptor that shares regions of expression with *period* and is well described also on the sub-cellular level for *P.dumerilii* (Poehn et al., 2022; Zurl et al., 2022). In line with the published expression patterns, the acquisition of SHInE-processed heads revealed expression of L-Cry in the medial brain nuclei and in the eyes (Figure 2C,D), in the same region as *period* RNA (Figure 2A,D).

### Co-visualisation of acetylated alpha-tubulin, *hox3* mRNA and EdU in posterior blastemas

Regenerating tissue after caudal amputation has been used in *Platynereis* for various studies of cell identity, regeneration and the hormonal control of maturation (Balavoine, 2015; Özpolat et al., 2021a; Özpolat and Bely, 2016; Planques et al., 2018; Rosa et al., 2005; Starunov et al., 2015). A key gene involved in both posterior growth as well as regeneration is *hox3*. It is expressed in a distinct set of cells believed to be ectoteloblasts (ectodermal stem cells). These cells are found in a region of high proliferation and expression of genes belonging to the germline multipotency program, called the segment addition zone (Gazave et al., 2013; Planques et al., 2018). We chose this gene for testing our protocol as it has a highly distinct, well-studied expression pattern. Cells positive for *hox3* also exhibit enlarged nucleoli, which makes it possible to identify them and validate the *hox3* staining.

Detection of *hox3* using HCR probes revealed a distinct region along the border between the pygidium and the regenerating segments (Figure 3E,H). We co-labelled samples using antiacetylated tubulin antibodies, which are established to visualize neurites of the nervous system. Antibody signal was found along the ventral nerve cord (VNC) and extended laterally into the animal’s parapodia and posteriorly into the newly regenerated anal cirri (Figure 3G,H). Both *hox3* HCR and anti-acetylated tubulin labeling match the established patterns, demonstrating the compatibility of these two techniques with the SHInE protocol.

As for the head samples, we also performed an additional labeling of proliferative cells by pre-treating animals with EdU for 30 minutes just before fixation, and visualized EdU incorporated in cells undergoing S-phase using click chemistry. The pattern of proliferation is broad and matches previous observations of regenerating tissue after posterior amputation (Figure 3F,H) (Planques et al., 2018). Taken together, the analyses performed in both head and posterior regenerates show that SHInE allows for co-detection of nuclear EdU with both mRNA and protein.

### SHInE is compatible with tissue clearing

We have previously presented a novel method on tissue clearing and depigmentation named DEEP-Clear. This method is compatible with various labeling techniques, including immunohistochemistry, fluorescent proteins and EdU-labeling. Enhanced transparency and refractive index matching allow for high-resolution imaging of specimens up to the centimeter range (Pende et al., 2020). While DEEP-Clear was shown to allow for the detection of RNA using conventional riboprobes, the employed chemistry includes an alkaline solution, in which RNA is expected to be degraded over time. As *in situ* HCR only requires binding to short pieces of RNA, rather than alignment with longer mRNA molecules, we reasoned that after clearing, *in situ* HCR should yield comparable or even superior results when compared to riboprobe-based *in situ* hybridization. As a suitable test case for assessing the compatibility of SHInE with tissue clearing, we focused on the aforementioned expression domains of *Pladu-period* in the eyes and the oval-shaped domain (Figure 4 B-B’) of the posterior medial forebrain. Whereas worm eyes are strongly pigmented, and the brain is opaque, DEEP-Clear had been shown to improve visualization of signals in both tissues (Pende et al., 2020).

Indeed, a comparison of untreated samples (Figure 4B’) and samples pre-processed using DEEP-Clear (Figure 4B) revealed that tissue clearing was not only compatible with *in situ* HCR, but improved the signal noticeably. This is even more apparent in the negative control, for which we again used unrelated sponge HCR probes (see Materials and Methods) with the same amplifier as for *Pladu-period*. Tissue clearing (Figure 4A) significantly reduced the background signal (Figure 4A’) that is likely caused by autofluorescence in the strongly pigmented eyes.

The degree to which tissue clearing improves signal might depend on the type of fluorophore used: When we investigated the expression of the *pdp1* gene – that is expressed in a similar pattern – using a spectrally different HCR amplifier (B1 amplifier coupled to Alexa 546), tissue clearing did not improve HCR signal intensity as it did for the B2 amplifier coupled to Alexa 647) (see Supplementary Figure S1). As expected, for blastemal tissue that is neither pigmented nor particularly opaque, clearing did not improve the HCR signal notably (Figure 4C-D’).

In summary, we find that SHInE is compatible with tissue clearing using DEEP-Clear, which, depending on the type of tissue and the fluorophore used for labeling, can improve the HCR signal quality and reduce background signal.

## Discussion

Here we present a novel protocol that combines *in situ* HCR with immunohistochemistry and EdU labeling of proliferating cells. The simultaneous visualization of RNA and proteins in the same tissue, as well as the high sensitivity of HCR-mediated RNA visualization, make the protocol suitable for detailed studies on the level of organisms, tissues and cells.

As we demonstrate, it is possible to successfully apply SHInE using home-made reagents, with HCR probes designed with a free webtool, home-made buffers and custom antibodies. This serves to keep the protocol affordable and flexible to many biological questions. The combination of multiple labeling methods within the same steps also greatly reduces the time required compared to similar protocols, further increasing the accessibility of these kinds of experiments, or allowing researchers to sample more replicates or experimental conditions.

Additionally, SHInE gives the researcher full control over reagents, which opens the protocol up for future improvement or adaptation to other models and tissues. For example, dextran sulfate, which is part of the probe amplification buffer, has been reported to negatively impact some antibodies during IHC (Callahan et al., 1991), so changing the concentration of the reagent could help in such cases. At several stages of the protocol, we tested different HCR probe concentrations and amplification lengths and report our findings and suggestions in the main protocol (Supplementary File 1), providing a foundation for other researchers to adapt and modify the protocol to their requirements.

As we show, compatibility of SHInE with tissue clearing allows for improved HCR signal intensity in originally opaque and pigmented tissue. Autofluorescence in the eyes significantly impacts imaging and obscures real fluorescent labeling in *Platynereis dumerilii* heads. Not only do we provide evidence for the compatibility of DEEP-Clear with *in situ* HCR, but furthermore show an improvement in visualization and signal to noise ratio, depending on tissue type and spectral range of the used fluorophores.

We believe that this protocol, with its flexibility and ease of use, combined with low cost and customization opportunities, will be a valuable resource. Whereas we have focused on *Platynereis* as a model species, the broad applicability of *in situ* HCR and DEEP-Clear for various invertebrate and vertebrate species suggests that the combined protocol we present here can easily be adapted for other model systems, and thereby help to complement the powerful possibilities opened by scRNAseq in a variety of model species.

## Acknowledgements

The research has been funded by the European Research Council (ERC) under the European Community’s Horizon 2020 Program ERC grant 81995 (K.T.-R.), the Distinguished Professorship program of the Helmholtz Society (K.T.-R.), the University of Vienna research platforms Rhythms of Life (F.R. and K.T.-R.) and Single-Cell Regulation of Stem Cells (F.R.), Austrian Science Fund [Fonds zur Förderung der Wissenschaftlichen Forschung (FWF)] projects I2972 (F.R.) and SFB F78 (F.R. and K.T.-R.). A.W.S is a recipient of a DOC Fellowship of the Austrian Academy of Sciences at the Max Perutz Labs.

The authors would like to thank the members of the Tessmar- and Raible labs for fruitful discussions and support in developing and testing this protocol, the marine animal facility, and the Max Perutz Labs BioOptics facility. We would also like to acknowledge the Özpolat lab at WUSTL, particularly Duygu Özpolat and Ryan Null, for sharing protocols, experience and their probe maker tool with us.

---

**Supplementary Figure S1.**
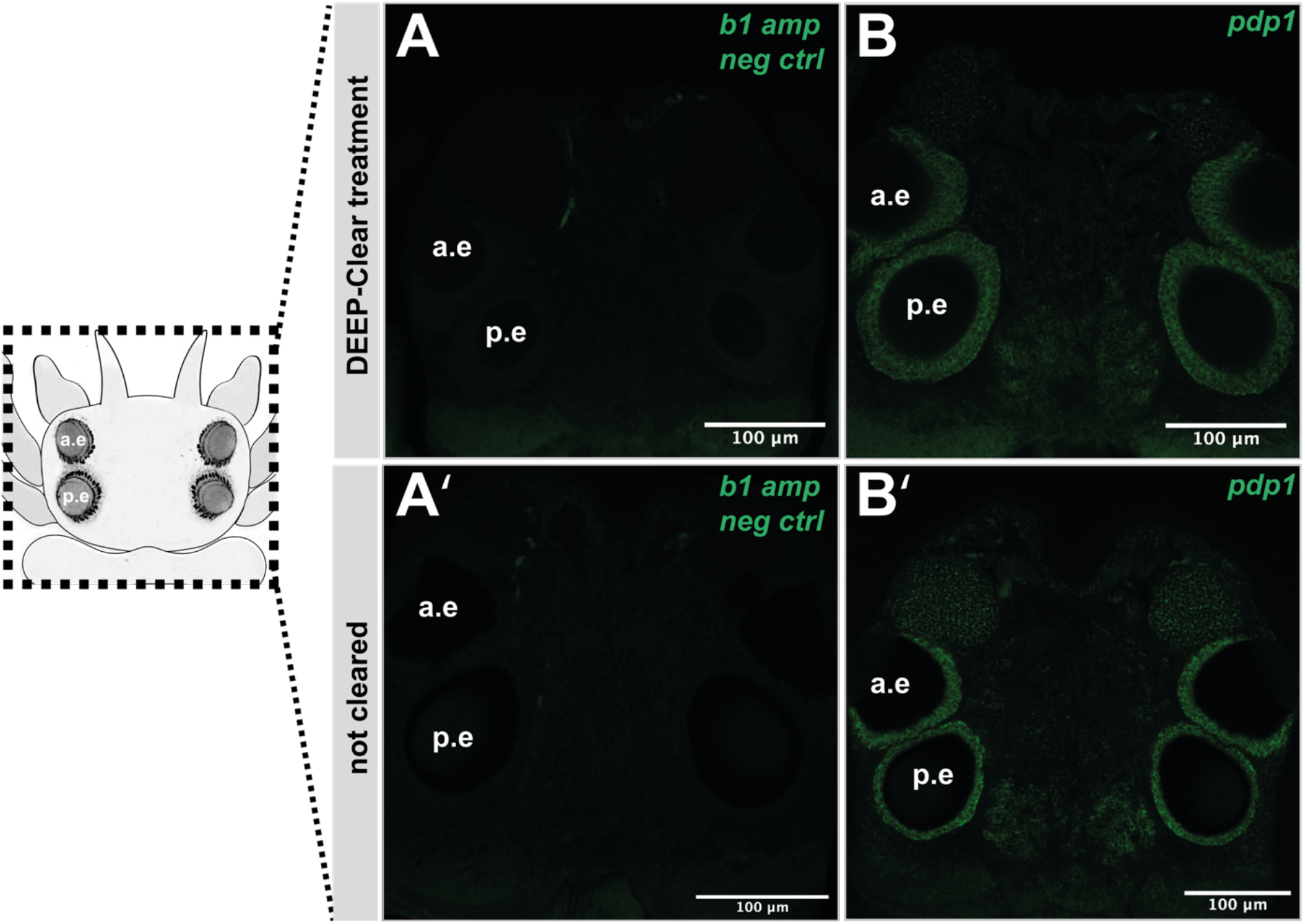
Assessment of the effects of tissue clearing on HCR signal intensity for detection using the Alexa-546-coupled B1 amplifier. Platynereis heads exhibit some autofluorescence around and in the eyes in the negative control of treated and untreated samples (A-A’). Pladu-pdp1 – detected using the Alexa-546-coupled B1 amplifier – is found around the eyes and nuclei of posterior medial forebrain. Following tissue clearing, HCR signal is particularly strong around the eyes. Less centralized Pladu-pdp1 is detected in both untreated and treated samples. For this detection, HCR signal seems more distinct and sharper in the untreated sample (B’) compared to the cleared specimen (B).

## References

Albayrak C, Jordi CA, Zechner C, Lin J, Bichsel CA, Khammash M, Tay S. 2016. Digital Quantification of Proteins and mRNA in Single Mammalian Cells. Mol Cell 61:914–924. doi:10.1016/j.molcel.2016.02.030

Balavoine G. 2015. Segment formation in Annelids: patterns, processes and evolution. Int J Dev Biol 58:469–483. doi:10.1387/ijdb.140148gb

Callahan LN, Phelan M, Mallinson M, Norcross and MA. 1991. Dextran Sulfate Blocks Antibody Binding to the Principal Neutralizing Domain of Human Immunodeficiency Virus Type 1 without Interfering with gpl20-CD4 Interactions. JOURNAL OF VIROLOGY 65, No.3:1543–1550.

Choi HMT, Calvert CR, Husain N, Huss D, Barsi JC, Deverman BE, Hunter RC, Kato M, Lee SM, Abelin ACT, Rosenthal AZ, Akbari OS, Li Y, Hay BA, Sternberg PW, Patterson PH, Davidson EH, Mazmanian SK, Prober DA, Rijn M van de, Leadbetter JR, Newman DK, Readhead C, Bronner ME, Wold B, Lansford R, Sauka-Spengler T, Fraser SE, Pierce NA. 2016. Mapping a multiplexed zoo of mRNA expression. Development 143:3632–3637. doi:10.1242/dev.140137

Choi HMT, Schwarzkopf M, Fornace ME, Acharya A, Artavanis G, Stegmaier J, Cunha A, Pierce NA. 2018. Third-generation in situ hybridization chain reaction: multiplexed, quantitative, sensitive, versatile, robust. Development 145:dev165753. doi:10.1242/dev.165753

Darmanis S, Gallant CJ, Marinescu VD, Niklasson M, Segerman A, Flamourakis G, Fredriksson S, Assarsson E, Lundberg M, Nelander S, Westermark B, Landegren U. 2015. Simultaneous Multiplexed Measurement of RNA and Proteins in Single Cells. Cell Reports 14:380–389. doi:10.1016/j.celrep.2015.12.021

Elagoz AM, Styfhals R, Maccuro S, Masin L, Moons L, Seuntjens E. 2022. Optimization of Whole Mount RNA multiplexed in situ Hybridization Chain Reaction with Immunohistochemistry, Clearing and Imaging to visualize octopus neurogenesis. Biorxiv 2022.02.24.481749. doi:10.1101/2022.02.24.481749

Fischer AH, Henrich T, Arendt D. 2010. The normal development of Platynereis dumerilii (Nereididae, Annelida). Front Zool 7:31. doi:10.1186/1742-9994-7-31

Fonseca RR da, Albrechtsen A, Themudo GE, Ramos-Madrigal J, Sibbesen JA, Maretty L, Zepeda-Mendoza ML, Campos PF, Heller R, Pereira RJ. 2016. Next-generation biology: Sequencing and data analysis approaches for non-model organisms. Mar Genom 30:3–13. doi:10.1016/j.margen.2016.04.012

Gazave E, Béhague J, Laplane L, Guillou A, Préau L, Demilly A, Balavoine G, Vervoort M. 2013. Posterior elongation in the annelid Platynereis dumerilii involves stem cells molecularly related to primordial germ cells. Dev Biol 382:246–267. doi:10.1016/j.ydbio.2013.07.013

Glossop NRJ, Hardin PE. 2002. Central and peripheral circadian oscillator mechanisms in flies and mammals. J Cell Sci 115:3369–3377. doi:10.1242/jcs.115.17.3369

Hauenschild C, Fischer A. 1969. Platynereis dumerilii. Mikroskopische Anatomie, Fortpflanzung, Entwicklung. Großes Zoologisches Praktikum.

Ibarra-García-Padilla R, Howard AGA, Singleton EW, Uribe RA. 2021. A protocol for whole-mount immunocoupled hybridization chain reaction (WICHCR) in zebrafish embryos and larvae. Star Protoc 2:100709. doi:10.1016/j.xpro.2021.100709

Kuehn E, Clausen DS, Null RW, Metzger BM, Willis AD, Özpolat BD. 2022. Segment number threshold determines juvenile onset of germline cluster expansion in Platynereis dumerilii. J Exp Zoology Part B Mol Dev Evol 338:225–240. doi:10.1002/jez.b.23100

Kuehn E, Stockinger AW, Girard J, Raible F, Özpolat BD. 2019. A scalable culturing system for the marine annelid Platynereis dumerilii. Plos One 14:e0226156. doi:10.1371/journal.pone.0226156

Özpolat BD, Bely AE. 2016. Developmental and molecular biology of annelid regeneration: a comparative review of recent studies. Curr Opin Genet Dev 40:144–153. doi:10.1016/j.gde.2016.07.010

Özpolat BD, Randel N, Williams EA, Bezares-Calderón LA, Andreatta G, Balavoine G, Bertucci PY, Ferrier DEK, Gambi MC, Gazave E, Handberg-Thorsager M, Hardege J, Hird C, Hsieh Y-W, Hui J, Mutemi KN, Schneider SQ, Simakov O, Vergara HM, Vervoort M, Jékely G, Tessmar-Raible K, Raible F, Arendt D. 2021a. The Nereid on the rise: Platynereis as a model system. Evodevo 12:10. doi:10.1186/s13227-021-00180-3

Özpolat BD, Randel N, Williams EA, Bezares-Calderón LA, Andreatta G, Balavoine G, Bertucci PY, Ferrier DEK, Gambi MC, Gazave E, Handberg-Thorsager M, Hardege J, Hird C, Hsieh Y-W, Hui J, Mutemi KN, Schneider SQ, Simakov O, Vergara HM, Vervoort M, Jékely G, Tessmar-Raible K, Raible F, Arendt D. 2021b. The Nereid on the rise: Platynereis as a model system. Evodevo 12:10. doi:10.1186/s13227-021-00180-3

Pende M, Vadiwala K, Schmidbaur H, Stockinger AW, Murawala P, Saghafi S, Dekens MPS, Becker K, Revilla-i-Domingo R, Papadopoulos S-C, Zurl M, Pasierbek P, Simakov O, Tanaka EM, Raible F, Dodt HU. 2020. A versatile depigmentation, clearing, and labeling method for exploring nervous system diversity. Sci Adv 6:eaba0365. doi:10.1126/sciadv.aba0365

Planques A, Malem J, Parapar J, Vervoort M, Gazave E. 2018. Morphological, cellular and molecular characterization of posterior regeneration in the marine annelid Platynereis dumerilii. Dev Biol 445:189–210. doi:10.1016/j.ydbio.2018.11.004

Poehn B, Krishnan S, Zurl M, Coric A, Rokvic D, Häfker NS, Jaenicke E, Arboleda E, Orel L, Raible F, Wolf E, Tessmar-Raible K. 2022. A Cryptochrome adopts distinct moon-and sunlight states and functions as sun-versus moonlight interpreter in monthly oscillator entrainment. Nat Commun 13:5220. doi:10.1038/s41467-022-32562-z

Raible F, Tessmar-Raible K. 2014. Platynereis dumerilii. Curr Biol 24:R676–R677. doi:10.1016/j.cub.2014.06.032

Rosa R, Prud’homme B, Balavoine G. 2005. caudal and even-skipped in the annelid Platynereis dumerilii and the ancestry of posterior growth. Evol Dev 7:574–587. doi:10.1111/j.1525-142x.2005.05061.x

Salic A, Mitchison TJ. 2008. A chemical method for fast and sensitive detection of DNA synthesis in vivo. Proc National Acad Sci 105:2415–2420. doi:10.1073/pnas.0712168105

Schindelin J, Arganda-Carreras I, Frise E, Kaynig V, Longair M, Pietzsch T, Preibisch S, Rueden C, Saalfeld S, Schmid B, Tinevez J-Y, White DJ, Hartenstein V, Eliceiri K, Tomancak P, Cardona A. 2012. Fiji: an opensource platform for biological-image analysis. Nat Methods 9:676–82. doi:10.1038/nmeth.2019

Schwarzkopf M, Liu MC, Schulte SJ, Ives R, Husain N, Choi HMT, Pierce NA. 2021. Hybridization chain reaction enables a unified approach to multiplexed, quantitative, high-resolution immunohistochemistry and in situ hybridization. Dev Camb Engl 148:dev199847. doi:10.1242/dev.199847

Stark R, Grzelak M, Hadfield J. 2019. RNA sequencing: the teenage years. Nat Rev Genet 20:631–656. doi:10.1038/s41576-019-0150-2

Starunov VV, Dray N, Belikova EV, Kerner P, Vervoort M, Balavoine G. 2015. A metameric origin for the annelid pygidium? Bmc Evol Biol 15:25. doi:10.1186/s12862-015-0299-z

Tainaka K, Kuno A, Kubota SI, Murakami T, Ueda HR. 2015. Chemical Principles in Tissue Clearing and Staining Protocols for Whole-Body Cell Profiling. Annu Rev Cell Dev Bi 32:1–29. doi:10.1146/annurev-cellbio-111315-125001

Tessmar-Raible K, Steinmetz PRH, Snyman H, Hassel M, Arendt D. 2005. Fluorescent two-color whole mount in situ hybridization in Platynereis dumerilii (Polychaeta, Annelida), an emerging marine molecular model for evolution and development. Biotechniques 39:460–464. doi:10.2144/000112023

Vieites-Prado A, Renier N. 2021. Tissue clearing and 3D imaging in developmental biology. Development 148:dev199369. doi:10.1242/dev.199369

Zantke J, Ishikawa-Fujiwara T, Arboleda E, Lohs C, Schipany K, Hallay N, Straw AD, Todo T, Tessmar-Raible K. 2013. Circadian and circalunar clock interactions in a marine annelid. Cell Reports 5:99–113. doi:10.1016/j.celrep.2013.08.031

Zattara EE, Özpolat BD. 2020. Developmental Biology of the Sea Urchin and Other Marine Invertebrates. Methods Mol Biology 2219:163–180. doi:10.1007/978-1-0716-0974-3_10

Zurl M, Poehn B, Rieger D, Krishnan S, Rokvic D, Rajan VBV, Gerrard E, Schlichting M, Orel L, Ćorić A, Lucas RJ, Wolf E, Helfrich-Förster C, Raible F, Tessmar-Raible K. 2022. Two light sensors decode moonlight versus sunlight to adjust a plastic circadian/circalunidian clock to moon phase. Proc National Acad Sci 119:e2115725119. doi:10.1073/pnas.2115725119

