## Supplementary File S1 for "A fast and versatile method for simultaneous HCR, immunohistochemistry and EdU labeling (SHInE)"

chn3t5gn

### SHInE - Simultaneous HCR, Immunohistochemistry, Nuclear staining and EdU V.(chn3t5gn)

2 Works for me

Reserved DOI:
10.17504/protocols.io.5qpvobnyzl4o/v1

Alexander Stockinger

DISCLAIMER

The research has been funded by the European Research Council (ERC) under the European Community’s Horizon 2020 Program ERC grant 81995 (K.T.-R.), the Distinguished Professorship program of the Helmholtz Society (K.T.-R.), the University of Vienna research platforms Rhythms of Life (F.R. and K.T.-R.) and Single-Cell Regulation of Stem Cells (F.R.), Austrian Science Fund [Fonds zur Förderung der Wissenschaftlichen Forschung (FWF)] projects I2972 (F.R.) and F78 (F.R. and K.T.-R.).A.W.S is a recipient of a DOC Fellowship of the Austrian Academy of Sciences at the Max Perutz Labs.

The authors would like to thank the members of the Tessmar- and Raible labs for fruitful discussions and support in developing and testing this protocol, our marine animal facility, and the Max Perutz Labs BioOptics facility. We would also like to acknowledge the Özpolat lab at WUSTL, particularly Duygu Özpolat and Ryan Null, for sharing protocols, experience and their probe maker tool with us.

ABSTRACT

This protocol allows the multiplexed use of four different molecular labelling techniques in whole-mount *Platynereis* tissues. In short, gene expression (via in situ HCR 3.0), cell proliferation (via EdU labelling), proteins (via Immunohistochemistry) and nuclei (DAPI/Hoechst staining) can be assessed in whole-mount tissue samples, allowing the co-visualisation of various types of molecules in the same specimen.

This protocol combines knowledge from multiple sources (see below), and is being submitted in parallel with a manuscript detailing the applications of the method. (We will add the reference as soon as it's available.)

We successfully applied this protocol to heads and posterior regenerates of the bristleworm *Platynereis dumerilii*, at various developmental stages of the animal. Given the general nature of the detected molecules, and the wide-spread use of the individual detection techniques, we anticipate that this protocol will be well applicable to a wider range of model systems.

References:

1) Choi HMT, Calvert CR, Husain N, Huss D, Barsi JC, Deverman BE, Hunter RC, Kato M, Lee SM, Abelin ACT, Rosenthal AZ, Akbari OS, Li Y, Hay BA, Sternberg PW, Patterson PH, Davidson EH, Mazmanian SK, Prober DA, Rijn M van de, Leadbetter JR, Newman DK, Readhead C, Bronner ME, Wold B, Lansford R, Sauka-Spengler T, Fraser SE, Pierce NA. 2016. Mapping a multiplexed zoo of mRNA expression. *Development* 143:3632–3637. doi:10.1242/dev.140137

2) Choi HMT, Schwarzkopf M, Fornace ME, Acharya A, Artavanis G, Stegmaier J, Cunha A, Pierce NA. 2018. Third-generation in situ hybridization chain reaction: multiplexed, quantitative, sensitive, versatile, robust. *Development* 145:dev165753. doi:10.1242/dev.165753

3) Kuehn E, Clausen DS, Null RW, Metzger BM, Willis AD, Özpolat BD. 2021. Segment number threshold determines juvenile onset of germline cluster proliferation in Platynereis dumerilii. *Biorxiv* 2021.04.22.439825. doi:10.1101/2021.04.22.439825

4) Pende M, Vadiwala K, Schmidbaur H, Stockinger AW, Murawala P, Saghafi S, Dekens MPS, Becker K, Revilla-i-Domingo R, Papadopoulos S-C, Zurl M, Pasierbek P, Simakov O, Tanaka EM, Raible F, Dodt H-U. 2020. A versatile depigmentation, clearing, and labeling method for exploring nervous system diversity. *Sci Adv* 6:eaba0365. doi:10.1126/sciadv.aba0365

5) Salic A, Mitchison TJ. 2008. A chemical method for fast and sensitive detection of DNA synthesis in vivo. *Proc National Acad Sci* 105:2415–2420. doi:10.1073/pnas.0712168105

6) Tessmar-Raible K, Steinmetz PRH, Snyman H, Hassel M, Arendt D. 2005. Fluorescent two-color whole mount in situ hybridization in Platynereis dumerilii (Polychaeta, Annelida), an emerging marine molecular model for evolution and development. *Biotechniques* 39:460–464. doi:10.2144/000112023

PROTOCOL INFO

Aida Coric, Alexander W. Stockinger, Petra Schaffer, Dunja Rokvic, Kristin Tessmar-Raible, Florian Raible . SHInE - Simultaneous HCR, Immunohistochemistry, Nuclear staining and EdU. **protocols.io**
https://protocols.io/view/shine-simultaneous-hcr-immunohistochemistry-nuclea-chn3t5gn
Version created by Alexander Stockinger

MANUSCRIPT CITATION please remember to cite the following publication along with this protocol

In submission

KEYWORDS

HCR, immunohistochemistry, in situ hybridisation, FISH, whole-mount, multiplexed, EdU, proliferation, Platynereis, Annelid

CREATED

Oct 10, 2022

LAST MODIFIED

Oct 12, 2022

PROTOCOL INTEGER ID

71099

GUIDELINES

- Please use aliquots of common stocks, including all buffers, to avoid repeated freeze-thaw cycles.
- The protocol is based on the detection of RNA, so minimize RNA degradation. It does not seem to be as critical as with sequencing experiments, but keep samples on ice whenever indicated, and work quickly in a clean setting.
- Protect samples from light once fluorescent dyes have been added to them!
- HCR signal seems to be best in the first 5 days after preparation of the samples, so ideally, image your samples right after preparing them.
- Many of the steps in this protocol can be optimized for individual samples or experimental conditions. We tried to indicate these steps. If you do come up with improved conditions for certain samples, we want to encourage you to upload a modified version of this protocol!

For multi-color fluorescence labeling, check whether the microscope you intend to use has lasers and filters available. HCR signal in general seems to only be visible on the confocal microscope. You can also check for bleed through risk and plan accordingly, for example using this tool:
https://www.thermofisher.com/order/fluorescence-spectraviewer

Probe maker:
Probes can be ordered from Molecular Instruments, alongside the amplifier hairpins that contain the fluorophore.
To design HCR probes yourself, you can use a Python based tool developed by Ryan Null in the Özpolat lab at WUSTL. It can be found here:
https://github.com/rwnull/insitu_probe_generator
and was first referenced in
Kuehn E, Clausen DS, Null RW, Metzger BM, Willis AD, Özpolat BD. 2022. Segment number threshold determines juvenile onset of germline cluster expansion in Platynereis dumerilii. *J Exp Zoology Part B Mol Dev Evol* 338:225–240. doi:10.1002/jez.b.23100

In short, use a target sequence and the probe generator will generate sets of probes that contain the initiator sequence of your choice.

* Probes can either be acquired through https://www.molecularinstruments.com, or self-designed (we use this python tool https://github.com/rwnull/insitu_probe_generator) and then ordered as oligos, for example through https://www.idtdna.com/pages/products/custom-dna-rna/dna-oligos/custom-dna-oligos/opools-oligo-pools

For more detailed instructions on EdU pulse-and-chase experiments in *Platynereis dumerilii*, please refer to
Zattara, E. E. & Özpolat, B. D. Developmental Biology of the Sea Urchin and Other Marine Invertebrates. *Methods Mol Biology* **2219**, 163–180 (2020).

MATERIALS TEXT

**Buffer recipes**

**Hybridization buffer (toxic, store at -20° C):**

| 1x concentration | For 40ml |
|---|---|
| 30% formamide | 12 mL formamide |
| 5x sodium chloride sodium citrate (SSC) | 10 mL of 20× SSC |
| 9 mM citric acid (pH 6.0) | 360 µL 1 M citric acid, pH 6.0 |
| 0.1% Tween 20 | 400 µL of 10% Tween 20 |
| 50 µg/mL heparin | 200 µL of 10 mg/mL heparin |
| 1x Denhardt's solution | 800 µL of 50× Denhardt's solution |
| 10% dextran sulfate | 8 mL of 50% dextran sulfate |
| | Fill to 40 ml with ultrapure H2O |

#### Wash buffer (toxic, store at -20° C):

| 1x concentration | For 40ml |
|---|---|
| 30% formamide | 12 mL formamide |
| 5× sodium chloride sodium citrate (SSC) | 10 mL of 20× SSC |
| 9 mM citric acid (pH 6.0) | 360 µL 1 M citric acid, pH 6.0 |
| 0.1% Tween 20 | 400 µL of 10% Tween 20 |
| 50 µg/mL heparin | 200 µL of 10 mg/mL heparin |
| | Fill up to 40 ml with ultrapure H2O |

#### Amplification buffer (store at 4° C):

| 1x concentration | For 40ml |
|---|---|
| 5× sodium chloride sodium citrate (SSC) | 10 mL of 20× SSC |
| 0.1% Tween 20 | 400 µl of 10% Tween 20 |
| 10% dextran sulfate | 8 mL of 50% dextran sulfate |
| | Fill up to 40 ml with ultrapure H2O |

#### 5x SSCT (store at 4° C):

| 1x concentration | For 40ml |
|---|---|
| 5× sodium chloride sodium citrate (SSC) | 10 mL of 20× SSC |
| 0,1% Tween 20 | 400µl of 10% Tween 20 |
| | Fill up to 40µl with ultrapure H2O |

**1X PTW:**

1X PBS with 0.1% Tween-20

**Tissue Clearing:**

1.Pende, M.*et al.* A versatile depigmentation, clearing, and labeling method for exploring nervous system diversity.*Sci Adv***6**, eaba0365 (2020).

**Reagents**

| A | B | C |
|---|---|---|
| **Reagent** | **Manufacturer** | **Product Number** |
| Citric acid monohydrate | Sigma-Aldrich | C1909 |
| Click-iT™ EdU Cell Proliferation Kit for Imaging, Alexa Fluor™ 488 dye | Invitrogen™ | C10337 |
| Denhardt's solution 50x | Invitrogen™ | 750018 |
| Dextran Sulfate 50% solution | Merck Millipore | S4030 |
| Formamide | Sigma-Aldrich | 47671 |
| Glycine | Roth | 3908.3 |
| Heparin | Sigma-Aldrich | H3393 |
| Hoechst 33342, Trihydrochloride, Trihydrate - 10 mg/mL Solution in Water | Invitrogen™ | H3570 |
| Magnesiumchlorid hexahydrate ($MgCl_2$) | VWR | 25.108.295 |
| Methanol | VWR | 20847 |
| Methanol, suitable for HPLC | Sigma-Aldrich | 34860 |
| Paraformaldehyde (PFA) | Sigma-Aldrich | 441244 |
| Phosphate buffered saline (PBS) | Sigma-Aldrich | P4417 |
| Proteinase K | Sigma-Aldrich | 1.245.680.100 |
| Saline-Sodium Citrate buffer 20× Concentrate (SSC 20X) | Sigma-Aldrich | S6639 |
| SlowFade™ Diamond Antifade Mountant | Molecular Probes™ | S36972 |
| Thymidine | Sigma-Aldrich | T9250 |
| Tween 20 | Sigma-Aldrich | P1379 |

DISCLAIMER:

The research has been funded by the European Research Council (ERC) under the European Community's Horizon 2020 Program ERC grant 81995 (K.T.-R.), the Distinguished Professorship program of the Helmholtz Society (K.T.-R.), the University of Vienna research platforms Rhythms of Life (F.R. and K.T.-R.) and Single-Cell Regulation of Stem Cells (F.R.), Austrian Science Fund [Fonds zur Förderung der Wissenschaftlichen Forschung (FWF)] projects I2972 (F.R.) and F78 (F.R. and K.T.-R.).A.W.S is a recipient of a DOC Fellowship of the Austrian Academy of Sciences at the Max Perutz Labs.

The authors would like to thank the members of the Tessmar- and Raible labs for fruitful discussions and support in developing and testing this protocol, our marine animal facility, and the Max Perutz Labs BioOptics facility. We would also like to acknowledge the Özpolat lab at WUSTL, particularly Duygu Özpolat and Ryan Null, for sharing protocols, experience and their probe maker tool with us.

Optional: EdU incubation

1

Incubate worms with [M]**10 micromolar (µM)** EdU in artificial sea water (ASW) (EdU aliquots in both DMSO and H2O work fine) for your desired pulse length.

2 1h

Optional: Chase out EdU with [M]**3 millimolar (mM)** thymidine for **01:00:00** ; then replace thymidine solution with fresh ASW and incubate for desired chase length

Day 1: Fixation, Dehydration

3 **Dissect tissue of interest**

Anesthetize worms in a 50% mix of 7.5% (w/v) MgCl2 and ASW

Dissect the tissue of interest; transfer it to a 1,5ml microcentrifuge tube pre-filled with 1ml ASW **On ice**

*Note: the protocol is also compatible with whole animals, but will require larger volumes and more reagent accordingly.*

4 1h

**Fix the samples**

Replace the ASW in the tube with 1ml 4%PFA/1XPTW (carefully pipet away the ASW; samples should sink to the tube bottom). Fix samples for **01:00:00** at **Room temperature** shaking gently on a rocking platform

*Make sure the samples are swimming back and forth freely in the tube; dislodge stuck samples by gently flicking the tube.*

5 11m

**Dehydrate and store samples**

Dehydrate the samples through a series of increasing MeOH concentrations diluted in 1X PTW.

Perform these steps on ice, and wait until the samples have settled down at the bottom of the well after each increase in MeOH concentration (takes around 1-3 minutes)

- 25% MeOH in PTW **00:03:00**

- 50% MeOH in PTW 00:03:00
- 75% MeOH in PTW 00:03:00
- Wash in100% MeOH 00:01:00
- Wash in100% MeOH 00:01:00

6

Store samples at -20° C for at least Overnight . Longer storage is possible (several months in our hands, but even longer storage might be possible).

Day 2: Rehydration, digestion, probe hybridization 1h 45m

7 12m

**Re-hydrate samples** through a series of decreasing MeOH concentrations in 1X PTW on ice.

- 75% MeOH in PTW 00:03:00
- 50% MeOH in PTW 00:03:00
- 25% MeOH in PTW 00:03:00
- Wash 2x in 1X PTW 00:03:00

8

**Perform proteinase K treatment** with 1ml of Proteinase K in PTW according to table below, at Room temperature .

8.1

This is a **critical, time-sensitive step** and all reagents needed for washes after digestion should be prepared in advance. Concentrations and digestion times **should be adjusted to individual tissues** and possibly differences in proteinase K batches.

| Tissue | PK concentration | time |
|---|---|---|
| Heads, Blastemas, adult tissues | 100µg/mL | 5' |
| Larvae, <1dpf | 100µg/mL | 30 seconds |
| Larvae, 3dpf | 100µg/mL | 2' |

8.2 Briefly rinse 2x with 1 ml of glycine wash buffer to stop digest. Work **On ice** from here on. 5m

Wash once for **00:05:00** with 1x PTW.

8.3 ⚠ 20m

Post-fix in 1mL 4%PFA diluted in 1xPTW for **00:20:00** shaking while still **On ice** .

8.4 Wash samples 2x **00:05:00** in 1ml PTW. 5m

9 ⊛ 20m

**Optional: tissue clearing**

If clearing with DEEP-CLEAR is wanted, perform it at this step. Worm tissue can be cleared at **37 °C** for **00:15:00** at 300 rpm on a heat block. Clearing time should be optimised for different tissues. Wash the samples in PTW thoroughly after clearing (3x **00:05:00** ).

*Note: in Platynereis tissues, the effects of clearing depend on the tissue sampled. Opaque tissues such as eyes benefit from clearing, while naturally transparent tissues like blastemas don't.*

*For details on tissue clearing, please refer to*

Pende, M.*et al.* A versatile depigmentation, clearing, and labeling method for exploring nervous system diversity.*Sci Adv***6**, eaba0365 (2020)

10 **Probe hybridization** with HCR probe(s) of choice.

*Note: a weak probe signal might be improved by increasing the probe concentration; several labs have reported successfully reusing the probe solution (simply freeze the probe mix after use)*

10.1 ⚠ 1h 5m

Incubate in 1ml 50% hybridization buffer in PTW for **00:05:00** at **Room temperature**

Incubate in 300µl 100% hybridization buffer for **01:00:00** at **37 °C** on a heat block .

*Careful when replacing media to not lose any samples. The buffer is very viscous; wait until samples settle down to the bottom of the tube.*

10.2

Meanwhile prepare the probe solution:

- 1pmol of each probe mix (1µl of 1µM stock) in 250µl hybridization buffer
- heat the probe solution to 37°C (for small volumes, we use the heat block the samples are already on; for larger volumes, we use a water bath)

10.3

Replace the hybridization buffer with the prepared probe solution and incubate the samples **Overnight** at **37 °C** .

Day 3: Washes 1h 16m 30s

11 1h 10m

**Wash out the HCR probes**:

Wash for **00:15:00** with 1ml pre-heated probe wash buffer at **37 °C**

Wash for **00:15:00** with 1ml pre-heated probe wash buffer at **37 °C**

Wash for **00:15:00** with 1ml pre-heated probe wash buffer at **37 °C**

Wash for **00:15:00** with 1ml pre-heated probe wash buffer at **37 °C**

Wash for **00:05:00** with 1ml 5X SSCT at **Room temperature**

Wash for **00:05:00** with 1ml 5X SSCT at **Room temperature**

**From here on, the protocol progresses differently for each combination of methods** (HCR only, HCR & EdU, HCR & IHC, HCR, EdU & IHC)
**Please select the step case below accordingly:**

Step 11 includes a Step case.

**HCR only**

**HCR & EdU**

**HCR & IHC**

**HCR, EdU & IHC**

Day 3: HCR probe detection 1h 16m 30s

step case

**HCR only**

This is the bare-bones protocol to just stain your samples with HCR.

12 **HCR Amplification:**

12.1 Wash with 1ml PTW for 00:05:00 at Room temperature 35m

Equilibrate samples in 300µl amplification buffer for 00:30:00 at Room temperature

12.2 **Meanwhile**: 31m 30s

Prepare the HCR hairpin mix.

- Use 15 pmol per hairpin (5µl of 3µM stock) - move them to individual 1,5ml microcentrifuge tubes*
- Heat the hairpins to 95 °C for 00:01:30 by adding them to a pre-heated heatblock
- Let the hairpins cool to Room temperature protected from light for 00:30:00
- Mix the cooled hairpins with amplification buffer to a total volume of 250µl

** NOTE: for each amplifier, TWO different hairpins have to be used! (e.g., B1H1 and B1H2 are a pair of hairpins compatible with the B1 amplifier sequence) - heat those in separate tubes and pool only directly before adding them to the sample!*

** NOTE: several probe pairs can be co-detected in one sample by using different sets of amplifiers with different fluorophores.*

12.3 Incubate the samples in 250 µl hairpin mix Overnight protected from light at Room temperature .

*Note: amplification buffer is viscous; samples will eventually equilibrate and sink to the bottom of the tube, but careful pipetting is recommended to not lose any samples.*

#### Day 4: Amplification Termination 1h 25m

13 **Terminate the amplification:** 1h 40m

Wash in 1ml 5X SSCT for 00:05:00 at Room temperature

Wash in 1ml 5X SSCT for 00:05:00 at Room temperature

Wash in 1ml 5X SSCT for 01:00:00 at Room temperature

Wash in 1ml 5X SSCT for 00:05:00 at Room temperature

Wash in 1ml PTW for 00:05:00 at Room temperature shaking on rocking platform

Staining of nuclei (DAPI/ Hoechst) can be performed here by adding the dye directly to PTW and increasing the incubation time to 00:15:00 .

Wash in 1ml PTW for 00:05:00 at Room temperature shaking on rocking platform

14 **Mount the samples:**
Mounting strategies may differ based on sample type/ size.
For whole mount *Platynereis* blastemas or heads, the following works well:

Equilibrate the sample in a small volume of mounting medium (in our hands, SlowFade Diamond worked best)
Put four small pieces of "pattafix" or similar poster mounting clay (Blu-tack or similar) into the corners of the "mounting area" of a microscope slide to build a stage.
Put the sample in the middle of the stage, and add a small volume (~10µl) of mounting medium on top of it.

Gently press down a cover glass, using a second microscope slide to keep it even.

#### Day 3: EdU click chemistry; HCR probe detection 1h 16m 30s

step case

##### HCR & EdU

Combines the HCR with EdU click detection.

12 **EdU detection with click chemistry:**

12.1 Wash samples in PTW for 00:05:00 at Room temperature (shaking on rocking platform) 10m
Wash samples in PTW for 00:05:00 at Room temperature (shaking on rocking platform)

12.2 Meanwhile, prepare the "click cocktail" according to manufacturer's protocol (see below):

*Note on reaction volume: depending on sample size, this could be enough for 1-2 1,5ml tubes; the staining reaction is extremely robust and strong in our hands, so a small volume (covering the samples fully) should be sufficient.*

For 500µl click cocktail, mix (in order):
- 430µl 1X reaction buffer
- 20µl $CuSo_4$*
- 1,2µl Alexa Fluor Azide
- 50µl reaction buffer additive (*Note: this is stored at 10X concentration! Dilute directly prior to use*)

** for samples with endogenous fluorescence, copper ions might be problematic. There is an alternative click reaction kit called "click-it plus" which works well with these samples and is also compatible with SHINE.*

12.3 Incubate with click-cocktail for 00:30:00 at Room temperature protected from light. 30m

#### 13 HCR Amplification:

13.1 Wash with 1ml PTW for 00:05:00 at Room temperature 45m

Wash with 1ml PTW for 00:05:00 at Room temperature

Wash with 1ml PTW for 00:05:00 at Room temperature

Equilibrate samples in 300µl amplification buffer for 00:30:00 at Room temperature

13.2 **Meanwhile**: 31m 30s

Prepare the HCR hairpin mix.

- Use 15 pmol per hairpin (~5µl of 3µM stock) - move them to individual 1,5ml microcentrifuge tubes*
- Heat the hairpins to 95 °C for 00:01:30 by adding them to a pre-heated heatblock
- Let the hairpins cool to Room temperature protected from light for 00:30:00
- Mix the cooled hairpins with amplification buffer to a total volume of 250µl

** NOTE: for each amplifier, TWO different hairpins have to be used! (e.g., B1H1 and B1H2 are a pair of hairpins compatible with the B1 amplifier sequence) - heat those in separate tubes and pool only directly before adding them to the sample!*

13.3

Incubate the samples in hairpin mix Overnight protected from light at Room temperature

Day 4: Amplification Termination; mounting 1h 25m

#### 14 Wash out the amplification mix: 1h 40m

Wash in 1ml 5X SSCT for 00:05:00 at Room temperature

Wash in 1ml 5X SSCT for 00:05:00 at Room temperature

Wash in 1ml 5X SSCT for 01:00:00 at Room temperature

Wash in 1ml 5X SSCT for 00:05:00 at Room temperature

Wash in 1ml PTW for 00:05:00 at Room temperature shaking on rocking

platform
Staining of nuclei (DAPI/ Hoechst) can be performed here by adding the dye directly to PTW and increasing the incubation time to 00:15:00 . Hoechst 33342 at a concentration of 10µg/ml works well in our samples.
Wash in 1ml PTW for 00:05:00 at Room temperature shaking on rocking platform

15 **Mount the samples:**
Mounting strategies may differ based on sample type/ size.
For whole mount Platynereis blastemas or heads, the following works well:

Equilibrate the sample in a small volume of mounting medium (in our hands, SlowFade Diamond worked best)
Put four small pieces of "pattafix" or similar poster mounting clay into the corners of the "mounting area" of a microscope slide to build a stage.
Put the sample in the middle of the stage, and add a small volume (~10µl) of mounting medium on top of it.

Gently press down a cover glass, using a second microscope slide to keep it even (see picture below).

Day 3: HCR probe detection; primary antibody 1h 16m 30s

step case

#### HCR & IHC

Combines HCR with immunolabelling

12 **HCR amplification & primary antibody:**

12.1 Wash with 1ml PTW for 00:05:00 at Room temperature (40m)
Wash with 1ml PTW for 00:05:00 at Room temperature
Equilibrate samples in 300µl amplification buffer for 00:30:00 at Room temperature

12.2 **Meanwhile**: (31m 30s)
Prepare the HCR hairpin mix.
- Use 15 pmol per hairpin (~5µl of 3µM stock) - move them to individual 1,5ml microcentrifuge tubes*
- Heat the hairpins to 95 °C for 00:01:30 by adding them to a pre-heated heatblock
- Let the hairpins cool to Room temperature protected from light for 00:30:00
- Mix the cooled hairpins with amplification buffer to a total volume of 250µl

** NOTE: for each amplifier, TWO different hairpins have to be used! (e.g., B1H1* *and B1H2 are a pair of hairpins compatible with the B1 amplifier sequence) - heat those in separate tubes and pool only directly before adding them to the sample!*

12.3 Add the primary antibody to the cooled and pooled hairpin mix (after addition of the amplification buffer).

*Note:*
*In our hands, a primary antibody dilution of 1:200 for commercial monoclonal antibodies worked well.*
*The dextrane sulfate in the amplification buffer can cause binding issues with antibodies. Adjusting its concentration could help, but was not necessary for the antibodies we used.*

12.4

Incubate the samples in hairpin + antibody mix **Overnight** protected from light at **Room temperature**

Day 4: Amplification Termination; secondary antibody 1h 25m

13 1h 25m

Wash in 1ml 5X SSCT for **00:05:00** at **Room temperature**
Wash in 1ml 5X SSCT for **00:05:00** at **Room temperature**
Wash in 1ml 5X SSCT for **01:00:00** at **Room temperature**
Wash in 1ml 5X SSCT for **00:05:00** at **Room temperature**

Wash in 1ml PTW for **00:05:00** at **Room temperature** shaking on rocking platform
Wash in 1ml PTW for **00:05:00** at **Room temperature** shaking on rocking platform

14

**Incubate with secondary antibody:**
Incubate in secondary antibody (500µl, 1:500 dilution) in PTW **Overnight** at **4 °C**

*Note: The ideal antibody concentration might be sample- and antibody dependent.*

14.1

Staining of nuclei (DAPI/Hoechst) can be performed during this step.
Hoechst 33342 at a concentration of 10µg/ml works well in our samples.

Day 5: Washes; Mounting 30m

30m

15 **Wash out secondary antibody:**

Wash in 1ml PTW for 00:15:00 at Room temperature shaking on rocking platform

Wash in 1ml PTW for 00:15:00 at Room temperature shaking on rocking platform

16 **Mount the samples:**

Mounting strategies may differ based on sample type/ size.
For whole mount Platynereis blastemas or heads, the following works well:

Equilibrate the sample in a small volume of mounting medium (in our hands, SlowFade Diamond worked best)
Put four small pieces of "pattafix" or similar poster mounting clay into the corners of the "mounting area" of a microscope slide to build a stage.
Put the sample in the middle of the stage, and add a small volume (~10µl) of mounting medium on top of it.

Gently press down a cover glass, using a second microscope slide to keep it even (see picture below).

Day 3: EdU click reaction; HCR probe detection; primary antibody 1h 16m 30s

step case

#### HCR, EdU & IHC

Combines all three techniques (EdU, HCR, IHC) into the full SHINE protocol.

12 **EdU decection with Click Chemistry:**

10m

12.1 Wash samples in PTW for 00:05:00 at Room temperature (shaking on rocking platform)

Wash samples in PTW for 00:05:00 at Room temperature (shaking on rocking platform)

12.2 **Meanwhile**, prepare the "click cocktail" according to manufacturer's protocol (see below):

*Note on reaction volume: depending on sample size, this could be enough for 1-2 1,5ml tubes; the staining reaction is extremely robust and strong in our hands, so a small volume (covering the samples fully) should be sufficient.*

For 500µl click cocktail, mix (in order):

- 430µl 1X reaction buffer
- 20µl $CuSo_4$*
- 1,2µl Alexa Fluor Azide
- 50µl reaction buffer additive (*Note: this is stored at 10X concentration! Dilute directly prior to use*)

** for samples with endogenous fluorescence, copper ions might be problematic. There is an alternative click reaction kit called "click-it plus" which works well with these samples and is also compatible with SHINE.*

12.3 Incubate with click cocktail for **00:30:00** at **Room temperature** protected from light. 30m

#### 13 HCR amplification & primary antibody:

13.1 Wash with 1ml PTW for **00:05:00** at **Room temperature** 45m

Wash with 1ml PTW for **00:05:00** at **Room temperature**

Wash with 1ml PTW for **00:05:00** at **Room temperature**

Equilibrate samples in 300µl amplification buffer for **00:30:00** at **Room temperature**

13.2 **Meanwhile**: 31m 30s

Prepare the HCR hairpin mix.

- Use 15 pmol per hairpin (~5µl of 3µM stock) - move them to individual 1,5ml microcentrifuge tubes*
- Heat the hairpins to **95 °C** for **00:01:30** by adding them to a pre-heated heatblock
- Let the hairpins cool to **Room temperature** protected from light for **00:30:00**
- Mix the cooled hairpins with amplification buffer to a total volume of 250µl

** NOTE: for each amplifier, TWO different hairpins have to be used! (e.g., B1H1 and B1H2 are a pair of hairpins compatible with the B1 amplifier sequence) - heat those in separate tubes and pool only directly before adding them to the sample!*

13.3 Add the primary antibody to the cooled and pooled hairpin mix (after addition of the amplification buffer).

*Note:*
*In our hands, a primary antibody dilution of 1:200 for commercial monoclonal antibodies worked well.*
*The dextrane sulfate in the amplification buffer can cause binding issues with antibodies. Adjusting its concentration could help, but was not necessary for the antibodies we used.*

13.4

Incubate the samples in hairpin + antibody mix **Overnight** protected from light at **Room temperature**

Day 4: Amplification termination; secondary Antibody; Mounting 1h 25m

14 **Terminate the amplification:** 1h 25m

Wash in 1ml 5X SSCT for **00:05:00** at **Room temperature**

Wash in 1ml 5X SSCT for **00:05:00** at **Room temperature**

Wash in 1ml 5X SSCT for **01:00:00** at **Room temperature**

Wash in 1ml 5X SSCT for **00:05:00** at **Room temperature**

Wash in 1ml PTW for **00:05:00** at **Room temperature** shaking on rocking platform

Wash in 1ml PTW for **00:05:00** at **Room temperature** shaking on rocking platform

15

**Incubate with secondary antibody:**

Incubate in secondary antibody (500µl, 1:500 dilution) in PTW **Overnight** at **4 °C**

*Note: The ideal antibody concentration might be sample- and antibody dependent.*

15.1

Staining of nuclei (DAPI/Hoechst) can be performed during this step.
Hoechst 33342 at a concentration of 10 µg/ml works well in our samples.

Day 5: Washes; mounting 30m

16 **Wash out secondary antibody:** 30m

Wash in 1ml PTW for **00:15:00** at **Room temperature** shaking on rocking platform

Wash in 1ml PTW for **00:15:00** at **Room temperature** shaking on rocking platform

17 **Mount the samples:**

Mounting strategies may differ based on sample type/ size.
For whole mount Platynereis blastemas or heads, the following works well:

Equilibrate the sample in a small volume of mounting medium (in our hands, SlowFade Diamond worked best)
Put four small pieces of "pattafix" or similar poster mounting clay into the corners of the "mounting area" of a microscope slide to build a stage.
Put the sample in the middle of the stage, and add a small volume (~10µl) of mounting medium on top of it.

Gently press down a cover glass, using a second microscope slide to keep it even (see picture below).
