## Supplementary File S2 for "A fast and versatile method for simultaneous HCR, immunohistochemistry and EdU labeling (SHInE)"

Supplementary File 2: HCR Probe sequences

A) Probe set for *Platynereis hox3* with B2 adapters *(Pladu_hox3*_b2)

CCTCGTAAATCCTCATCAaaAATGTTGTTGCGGGTTGTTGTGTGG
GTTGGATGTGAGGATGTTGCTGTCGaaATCATCCAGTAAACCGCC
CCTCGTAAATCCTCATCAaaTTGCTGGTGAGCTTGACTATGCGGA
ATGAGGCATTGGCTGTTGTAAATGCaaATCATCCAGTAAACCGCC
CCTCGTAAATCCTCATCAaaTGTTGATTATGGTGCTGCTGCATAT
GGTTGGGTATGCTGACTGTGAGGATaaATCATCCAGTAAACCGCC
CCTCGTAAATCCTCATCAaaGTGTATTGGCGGAAGGAGGAGTTTC
GCTCCTGATGCATGTTCGGAGGCATaaATCATCCAGTAAACCGCC
CCTCGTAAATCCTCATCAaaTAGGGGGACACCCTTCTCACTCGGA
AGGGCCGGACATCGGTCCAGAATGAaaATCATCCAGTAAACCGCC
CCTCGTAAATCCTCATCAaaTCCCCATCTTCGCTCATCTCTTTTT
AGATCTTCCATACTCCTTCCTCCGTaaATCATCCAGTAAACCGCC
CCTCGTAAATCCTCATCAaaTGTACTTCATGCGTCTATTCTGGAA
AATTGGGCTTCAACCTCGAATCTTTaaATCATCCAGTAAACCGCC
CCTCGTAAATCCTCATCAaaCAACAGTGCTGCCATCTCTATTCTC
AATCTTAATCTGTCTCTCGCTCAAAaaATCATCCAGTAAACCGCC
CCTCGTAAATCCTCATCAaaAACTCCTTCTCCAACTCCACCAATT
GGCCGACACAGATATCTGTTGAAATaaATCATCCAGTAAACCGCC
CCTCGTAAATCCTCATCAaaTGCTCGGTTTCTCGCTCTCTCCGCT
CTGACGTATAAGCGGTTCTTGCTCTaaATCATCCAGTAAACCGCC
CCTCGTAAATCCTCATCAaaTCCGGTTCCTGGTCCGGCGCTCTCC
CGAGTCTCCTCCGTTGATGCTTCCAaaATCATCCAGTAAACCGCC
CCTCGTAAATCCTCATCAaaTTCCCTGGGTTGGGTCTCTTGTTGC
CTAATTCCATCATCTGATCCTGAGGaaATCATCCAGTAAACCGCC
CCTCGTAAATCCTCATCAaaCAGGGGTCAACCCCGGCCCCATGGA
GGGGGCCCTTCATCATCAGGGGTCCaaATCATCCAGTAAACCGCC
CCTCGTAAATCCTCATCAaaGTTGAATGTGTTGTAGTCCATAGGG
ACCAGGGATCCCATACTGCATACAGaaATCATCCAGTAAACCGCC
CCTCGTAAATCCTCATCAaaGCCATGTATTGGGGTCTGTAAGGAT
GAACAATGTTGGTTCATCTGACCATaaATCATCCAGTAAACCGCC
CCTCGTAAATCCTCATCAaaCCATATATCCCGGATAGCCATTTGT
GGGAGGGCTGATCCCCTCCATAGCTaaATCATCCAGTAAACCGCC
CCTCGTAAATCCTCATCAaaTTGGCTCTCGTAATAAGGTCCTTTG
TTGGTAAAATCCAGGATGGAATGAGaaATCATCCAGTAAACCGCC
CCTCGTAAATCCTCATCAaaACGGAGTTGACAGCACTGTAGGGGA
TTCTGAAGGTGCATAGGATCCCTCGaaATCATCCAGTAAACCGCC
CCTCGTAAATCCTCATCAaaACTTAACTTTCAACTGTCAATCAGA
CATTGATTGGTGCTGACATTGTGACaaATCATCCAGTAAACCGCC
CCTCGTAAATCCTCATCAaaATGTCAGCTGTATGAGTGTCTTAAA

TTAGCAAATCTAGGTACATATGAAGaaATCATCCAGTAAACCGCC
CCTCGTAAATCCTCATCAaaAGGCTTGTCAGAGAAGGTATATAGT
GCAGCCTCATACATGTATAGATGACaaATCATCCAGTAAACCGCC
CCTCGTAAATCCTCATCAaaACTGTCATTCCCAACTCATCTCTGG
ATGGTGATGTCTTGAGCTGGTGTGTaaATCATCCAGTAAACCGCC
CCTCGTAAATCCTCATCAaaCTCTCTTGAGAGGTCACTTCTATGA
CATGTCTTCTGTTCCGACTCTAGGAaaATCATCCAGTAAACCGCC
CCTCGTAAATCCTCATCAaaGAACCCGGACCTTGAAGAATGGCTC
CACACTACTCTCTGGAAGTCACACAaaATCATCCAGTAAACCGCC
CCTCGTAAATCCTCATCAaaCTTTGGATGGTCAGGCCTCACTTTT
AATCTCCTGGCTTCTGTCCTTCCTTaaATCATCCAGTAAACCGCC
CCTCGTAAATCCTCATCAaaGCCTCACTCTGGGATGGCCTGAGCC
TTTTACAGCCTCACTTTGGATTGCCaaATCATCCAGTAAACCGCC
CCTCGTAAATCCTCATCAaaGGAATAATGGCGTGAAATTGTTAAT
CTTTTGGCTTGATGTGTCTCAAATGaaATCATCCAGTAAACCGCC
CCTCGTAAATCCTCATCAaaGGCGATGATGCTCCCAATCATTGAT
TTTCTATCTCGCGTTGACTTCCATTaaATCATCCAGTAAACCGCC
CCTCGTAAATCCTCATCAaaAGTGTTGGAACCAATGTATATTGTG
ATCTTTAATTGTGTCCAATAAGTTTaaATCATCCAGTAAACCGCC
CCTCGTAAATCCTCATCAaaTTGGCAACGTTTTCCAACGAAAGAC
AAACTATGGTATTATGTTTTTGGTAaaATCATCCAGTAAACCGCC

B) Probe set for *Platynereis period* with B2 adapters *(Pladu_per*_b2)

CCTCGTAAATCCTCATCAaaGTCATCGCTCTTAAACAGCTGCTCC
TCAGTCCATTTCAACTGAATCAAAAaaATCATCCAGTAAACCGCC
CCTCGTAAATCCTCATCAaaGCCCTCGAACATTGAGGTCATTGAA
ATGTAAAACACTGTCCGAACATTCCaaATCATCCAGTAAACCGCC
CCTCGTAAATCCTCATCAaaGGTAATGGTGCTCGGGGAACACCGC
GGGGCTGTCGCTGCTAGACGAGCAGaaATCATCCAGTAAACCGCC
CCTCGTAAATCCTCATCAaaGTTAGTAAGCGGTTGCGGTTGTTTC
TACGCTCTTCAGTGCTGTGATCTGCaaATCATCCAGTAAACCGCC
CCTCGTAAATCCTCATCAaaCGACCCCTCCTGCATTGCTTCCTCT
ACTGGCTGTCGTTGACACCCCATCTaaATCATCCAGTAAACCGCC
CCTCGTAAATCCTCATCAaaGATTATGGTCTCGTTCGGTCGTGTC
GTCAAACCTTTCAGCGTCTTTATCAaaATCATCCAGTAAACCGCC
CCTCGTAAATCCTCATCAaaTAGCTCGGGCTGGGGGCAGTAATCA
GACGAGTTCATCCAACTGTTCTCTGaaATCATCCAGTAAACCGCC
CCTCGTAAATCCTCATCAaaAGGGACAAACAACCTCATGAGCATG
GGGCATGGCGACGAAATCTATCTTGaaATCATCCAGTAAACCGCC
CCTCGTAAATCCTCATCAaaGCTGGGAGTCAGGGAGCTCGATCCT
CGTCACCTCCTCGTGACTCCTCTGTaaATCATCCAGTAAACCGCC
CCTCGTAAATCCTCATCAaaGTCTTCCGCTCCCTCCTCGCTATCT

CCGTTCCTCCACAACCTGTTTATTCaaATCATCCAGTAAACCGCC
CCTCGTAAATCCTCATCAaaTCTTCCAGCTGGCTCATCAGTTCCT
TCAATGTCTTCCTCCTCAGGGGGCAaaATCATCCAGTAAACCGCC
CCTCGTAAATCCTCATCAaaGGTGGCGCTGCCGGCAGCAGCTTCT
TCCGACCAGTTCACCGCCGTGACCCaaATCATCCAGTAAACCGCC
CCTCGTAAATCCTCATCAaaTGGTTGCTCGTCTCGCCCGTTTTCA
AAACTCATCTCGTCCGACACTTCTTaaATCATCCAGTAAACCGCC
CCTCGTAAATCCTCATCAaaCAGCTTGGGGATGGTAAAACGGCCA
GGTAGAAGAAGGGCTGCATGATTGAaaATCATCCAGTAAACCGCC
CCTCGTAAATCCTCATCAaaTGCCTTGCTGTTGAGCACACATATC
TCACTGAACTCTGGATCATGCCGCTaaATCATCCAGTAAACCGCC
CCTCGTAAATCCTCATCAaaCGTCGTTCAACAACCTCTTCTTTCT
ATTTAGCCGCCTTCTTGTGACCAGCaaATCATCCAGTAAACCGCC
CCTCGTAAATCCTCATCAaaGGGGCTTGGGGGCATGACTTCCTCT
TGAATATTCGGGAACCTCCTCCTGGaaATCATCCAGTAAACCGCC
CCTCGTAAATCCTCATCAaaCTGTCCAACCTCCTCGAGGCCTTCT
AGCTTTAGCGTAAGGAGTTGCTTGCaaATCATCCAGTAAACCGCC
CCTCGTAAATCCTCATCAaaCTCCTTGGAGGACTCCTCTTTCGTT
TTGAGTGGGGCTTGCTCTTTCTTCTaaATCATCCAGTAAACCGCC
CCTCGTAAATCCTCATCAaaGGACATGGGGTAGGGTGGTGCGACG
CCCAGGGGTTGGCTCAAGGCTGGAGaaATCATCCAGTAAACCGCC
CCTCGTAAATCCTCATCAaaGAAGTCAAAACGTTTGGTCCAAGGG
CTGTACGACTTCATGTTGACCCATGaaATCATCCAGTAAACCGCC
CCTCGTAAATCCTCATCAaaCGGATGGTAATATTCGAACACTGAA
TATTCCAAGAAGCAGCTGCAGATCTaaATCATCCAGTAAACCGCC
CCTCGTAAATCCTCATCAaaCAAAGGCAAGCACTCTAAAACCAGG
TGAACAGGGCTTTGAATAAGCAGAAaaATCATCCAGTAAACCGCC
CCTCGTAAATCCTCATCAaaTCAGGCTCCTCGCCACTGGATTCAC
CGGAAGAAGAACATTGTCTTCCTCTaaATCATCCAGTAAACCGCC
CCTCGTAAATCCTCATCAaaCCAAGCACTTCTGACAAATCAGGGT
TGGTTGAACCAATAATTCTTAGGATaaATCATCCAGTAAACCGCC
CCTCGTAAATCCTCATCAaaGCTCCTCCCGATTCCTCCCCCTTGA
GCACCAAGGAGTTCTATGTCTGCTCaaATCATCCAGTAAACCGCC
CCTCGTAAATCCTCATCAaaTTGTCTGAGATGCCGCTACTGCATC
TTCTTCTTGTCCTTTCTGTTTTGTGaaATCATCCAGTAAACCGCC

C) Probe set for *Platynereis pdp1* with B1 adapters *(Pladu_pdp1*_b1)

GAGGAGGGCAGCAAACGGaaTTACAAATCAGACAAAATACACCTA
TTTTTGAAAATGTCCATGCATTTTAtaGAAGAGTCTTCCTTTACG
GAGGAGGGCAGCAAACGGaaAGTGTAATATATATCCCATACACAC
GCAAATATATTACATTATTGATAATtaGAAGAGTCTTCCTTTACG
GAGGAGGGCAGCAAACGGaaAGCCTTGATTATGCATATATTCTTC

TTTCACGAGGCTCACTTGGTCACATtaGAAGAGTCTTCCTTTACG
GAGGAGGGCAGCAAACGGaaAAAGGTTACACCTGGATTTTATCAA
TATTAAGGAATGTACTACATTAATGtaGAAGAGTCTTCCTTTACG
GAGGAGGGCAGCAAACGGaaATATTTCTTGTCCGATATTAAGTTT
TCTCATAGACACATTTAGTCATGTTtaGAAGAGTCTTCCTTTACG
GAGGAGGGCAGCAAACGGaaTATTTCATTCACTTCTTAACAACAT
ATAGGTGCGTCTAATTTTGTTTCTTtaGAAGAGTCTTCCTTTACG
GAGGAGGGCAGCAAACGGaaTGGCATTAAACTTAAATATCAAGAT
ACTAACAGCGAACTATTGTAAGCAAtaGAAGAGTCTTCCTTTACG
GAGGAGGGCAGCAAACGGaaGTTAGTGAAAACAAAACATTGTTAA
TTTTTACTGACAAATCAACAAGTGTtaGAAGAGTCTTCCTTTACG
GAGGAGGGCAGCAAACGGaaGAACGTAAATTTAAGTACCACAAGT
CCATACATTGCATTCTCTCAAACAAtaGAAGAGTCTTCCTTTACG
GAGGAGGGCAGCAAACGGaaATGTAAAATAGTCAATAACGATATA
CGACAACATCTATTTACAGTGTACAtaGAAGAGTCTTCCTTTACG
GAGGAGGGCAGCAAACGGaaCCACCACCTTGCCGTAAAAACGCAT
TTACAATCTAACTGGATCTGTCCGGtaGAAGAGTCTTCCTTTACG
GAGGAGGGCAGCAAACGGaaCAACTCCCTGAGATTTTAACATCTA
CAGGTGACCACCATTAACTAACATCtaGAAGAGTCTTCCTTTACG
GAGGAGGGCAGCAAACGGaaTGCAACTGGAACAATCCAGTCCTTG
TAAATATTTGTTCACTCAATTCATAtaGAAGAGTCTTCCTTTACG
GAGGAGGGCAGCAAACGGaaATAAATCAGTAAGATTCATCAGTTT
TCAAAAGTCTAAGGCAAGATCGGATtaGAAGAGTCTTCCTTTACG
GAGGAGGGCAGCAAACGGaaTATTTGTTTTGAGTTAAATATCATA
TCATATCAAACCAATTACAGGTAAAtaGAAGAGTCTTCCTTTACG
GAGGAGGGCAGCAAACGGaaTTTAATTAACTGCACAGTGCGCAAT
AGCAGGAGATAAACCCTTAAATATTtaGAAGAGTCTTCCTTTACG
GAGGAGGGCAGCAAACGGaaTAAAGACTACCTGTCAATCGCCTAT
AGTGCTGCTACAATATAATTATATAtaGAAGAGTCTTCCTTTACG
GAGGAGGGCAGCAAACGGaaAGATAGCATATATACACTCAATTCT
ATTGACCATCAGAGAGGCCAGTGTCtaGAAGAGTCTTCCTTTACG
GAGGAGGGCAGCAAACGGaaACTTACAAAACCCATTATAAAGATT
GGTACACCATATGTAGACATTCGATtaGAAGAGTCTTCCTTTACG
GAGGAGGGCAGCAAACGGaaAAACCTATCACTCTCTGCTCCTAAT
GGAGATAAATTGTCATAAGAGACAGtaGAAGAGTCTTCCTTTACG
GAGGAGGGCAGCAAACGGaaCTGACAAGATATTTCCATCACTTGC
CAGATTTCTGAAGGTATCGGTTCTCtaGAAGAGTCTTCCTTTACG
GAGGAGGGCAGCAAACGGaaTACTGCTTTAGTACGATTTACTTCT
CCAATCATACTTTTACCCTCCCCTAtaGAAGAGTCTTCCTTTACG
GAGGAGGGCAGCAAACGGaaTTCAATTTTATCTTCAAGCAAAATT
TGAGATATTCCAAGTACTGTTTTAAtaGAAGAGTCTTCCTTTACG
GAGGAGGGCAGCAAACGGaaAACAGCCAACCACACTAAAATGCTT
TATAGCCTTTCATTCTGTTTAATAAtaGAAGAGTCTTCCTTTACG

GAGGAGGGCAGCAAACGGaaATGAAAACTTGTAATATGGCTGCTG
TTTATAGTAATGACAATTTTACGTCtaGAAGAGTCTTCCTTTACG
GAGGAGGGCAGCAAACGGaaGTATATAGCACAAGATACAACGATT
ATCCAATGTAGAAAAGAGCCATGGAtaGAAGAGTCTTCCTTTACG
GAGGAGGGCAGCAAACGGaaAATTGCATTGCCTCTTTGCTACCTT
ATATTTATGACGTCTAAGAATTATAtaGAAGAGTCTTCCTTTACG
GAGGAGGGCAGCAAACGGaaTTGAACTGTACATCATGGGGGAACT
CTTTTTGCAGGAAAATCATACAATAtaGAAGAGTCTTCCTTTACG
GAGGAGGGCAGCAAACGGaaGAGTGTGCTTAACCTGGCGCAGTGC
CGAGATGGAACTAATTACATTGGCTtaGAAGAGTCTTCCTTTACG
GAGGAGGGCAGCAAACGGaaGCACTTAGCACCAGGCAAGTACTTT
ACTTAAGGCATCCTCCGTGATGTAAtaGAAGAGTCTTCCTTTACG
GAGGAGGGCAGCAAACGGaaGCCGCTGTACGCCTCTCTTGAGGAC
ATAGTCTATACACCAATGGGTGCAAtaGAAGAGTCTTCCTTTACG
GAGGAGGGCAGCAAACGGaaAGCTGCTTGAGGGAACTTCCTCCTT
ATGACTGGCTCGGTAAGCTCGGGCTtaGAAGAGTCTTCCTTTACG
GAGGAGGGCAGCAAACGGaaTTCCAACTTAGCCATCTTTTTCTTC
TTCCACAGTCTTCTGACCGAGCATGtaGAAGAGTCTTCCTTTACG
GAGGAGGGCAGCAAACGGaaTTGTCGTCTCGCAGTCCGTGGTTTT
TCTTTCATTTCGAGACGCAGTTTCTtaGAAGAGTCTTCCTTTACG
GAGGAGGGCAGCAAACGGaaTCTGGTTCTCCTTGATGCGGCGTGC
TCTCCAAGAACGCTGCTCGGAGCGCtaGAAGAGTCTTCCTTTACG
GAGGAGGGCAGCAAACGGaaCTTCTTCCTGCGGCACCAGTACTTG
ACGTGACCTCTTTGCGGCAACGTTGtaGAAGAGTCTTCCTTTACG
GAGGAGGGCAGCAAACGGaaTTTCTGGACTTCTTGATCATAGGCT
TCCTTGAGATCTTCGGGTACAAAAAtaGAAGAGTCTTCCTTTACG
GAGGAGGGCAGCAAACGGaaTCTTTCTTGGATCGAAGCCTTCGTG
GCCGCAACTCCTCCAGACTGAAACTtaGAAGAGTCTTCCTTTACG
GAGGAGGGCAGCAAACGGaaGGGCCAGTGCTCGGGAACTTGGGCA
ATGGTTGTGGGAGCGGTTATATTCAtaGAAGAGTCTTCCTTTACG
GAGGAGGGCAGCAAACGGaaTCCACTTTGGGCGAACCGGGCCCTG
GAAAGGTCGGACTCTGAAACCATAAtaGAAGAGTCTTCCTTTACG
GAGGAGGGCAGCAAACGGaaCGGGCGAGGGTCCTTTCACAGGAGA
ATGAGTTTGAGTGGGTTGGGGAAGCtaGAAGAGTCTTCCTTTACG
GAGGAGGGCAGCAAACGGaaTTTAGTTGGGGACCCGGATGTGGAT
AGTGCGAGGGTCAACCATGACTTCTtaGAAGAGTCTTCCTTTACG
GAGGAGGGCAGCAAACGGaaGGAGATTTTTCTCCTTCCAGAGCCG
GCAAGCTGTTGACCGGGTGATACCGtaGAAGAGTCTTCCTTTACG
GAGGAGGGCAGCAAACGGaaTGCCATTCTCATGCAAGAACTCATC
CCTGAGCCTCCTGCATGGCTAGGGGtaGAAGAGTCTTCCTTTACG
GAGGAGGGCAGCAAACGGaaATAAGATTTGTCCCACAAGTTGGGG
GTCCATATACTCCAGTTTGAAGTCGtaGAAGAGTCTTCCTTTACG
GAGGAGGGCAGCAAACGGaaAATCCCAGGTCTTCCTCATCCTTCT

A fast and versatile method for simultaneous HCR, immunohistochemistry and EdU labeling (SHInE)

AGGAAAGCCCCCTGGCCGGGAAACAtaGAAGAGTCTTCCTTTACG
GAGGAGGGCAGCAAACGGaaTTGGAGGGTTGACTAGATTAGGATC
CTTCTTTGTTTCCGACTGTGGGTAGtaGAAGAGTCTTCCTTTACG
GAGGAGGGCAGCAAACGGaaTTGATAGAGCGACATTTTTCAGCGT
CAGAAGCGCCTTCAAAGTCATTCCGtaGAAGAGTCTTCCTTTACG
GAGGAGGGCAGCAAACGGaaTCCGTCAGTCCTGTTCTGTGTGCGT
ATTTCTCAAAACACTCTCGTCTTATtaGAAGAGTCTTCCTTTACG
